# Adipocytes are dispensable in shaping the ovarian cancer tumor microenvironment in the omentum

**DOI:** 10.1101/2025.10.28.685098

**Authors:** Rachel L. Mintz, Jichang Han, Emily G. Butka, Alexandre Gallerand, Ayoung Kim, Sarah Ning, Nicole K. H. Yiew, Mary Wohltmann, Wei Zou, Nan Zhang, Samantha A. Morris, Bernd H. Zinselmeyer, Gwendalyn J. Randolph

## Abstract

The omentum, a specialized adipose tissue within the peritoneum, is a primary niche for ovarian cancer (OC) dissemination during peritoneal carcinomatosis. Traditionally, omental adipocytes are thought to promote OC growth by supplying lipids, supported by evidence that global FABP4 deficiency reduces tumor progression. Here, we generated mice lacking mature adipocytes in the peritoneum, including the omentum. ID8p53^−/−^Brca2^−/−^, BPPNM, and KPCA OC cells retained a propensity to seed regions typically associated with adipocytes, even without mature adipocytes. However, the lack of mature adipocytes did not suppress peritoneal OC expansion, whereas removing the adipocyte-free omentum did. Murine and human single-cell RNA sequencing revealed that endothelial FABP4 was high in the omentum. Indeed, endothelial cell-selective deficiency of FABP4 reduced OC growth in the peritoneum. These findings prompt a reevaluation of adipocyte contributions to OC progression and suggest a key role of the omental vasculature in supporting OC metabolic growth.

## Introduction

At diagnosis, over 80% of ovarian cancer (OC) cases have metastasized to the peritoneal cavity, contributing to a five-year survival rate of only 25%^1^. The greater omentum, a highly vascularized, specialized visceral adipose tissue fold that arises off the greater curvature of the stomach, is an early and predominant site of transcoelomic metastasis in OC^2^. As cancer cells home to the omentum, tumor nodules eventually coalesce and replace the underlying normal fat, resulting in profound “omental caking.” Structurally, the human omentum is composed of two distinct regions, a vascularized, adipocyte-rich compartment and a translucent, mesothelial sheet, each with a unique cellular composition^3^. As tumor burden increases, these compartments undergo remodeling, with adipocytes replaced by fibroblasts, lymphocytes, and macrophages to varying extents, altering the underlying matrix structure^4^. Consequently, the omentum is removed in debulking surgeries as a standard of care for advanced disease, and selective omentectomies are debated for earlier disease stages^5,6^. Furthermore, omentectomy reduces the growth of experimental OC in the peritoneum of mice^7,8^. Still, it remains unknown why peritoneal metastases gravitate toward this specific omental niche.

Bolstered by the conclusions of a 2011 landmark study from the Lengyel group^9–11^, the notion that adipocytes drive OC peritoneal metastasis has prevailed^12,13^. In addition to secreting IL-8, IL-6, monocyte chemoattractant protein-1, and adiponectin, adipocytes can transfer lipids to OC tumor cells in vitro^9,14^. The concept that local lipid transfer provides metabolic support to tumors has been applied to other cancers^15–17^, particularly those in which white adipocytes are found near the organs containing the primary tumor, such as breast, prostate, colon, and skin cancer. The strongest mechanistic explanation for this metabolic relationship emerged from evidence that ovarian tumor cell expansion in the mouse peritoneal cavity was greatly suppressed in mice lacking fatty acid-binding protein 4 (FABP4), also known as adipocyte protein 2, in all host cells^9^. Given that FABP4 accounts for 1-3% of soluble protein in mature adipocytes^18^, this finding in FABP4 knockout mice, in conjunction with images of tumor cell-adipocyte proximity in patient samples, fueled the concept that fatty acids from host adipocytes support tumor seeding and progression.

The omentum in humans serves as the largest visceral fat depot. Accordingly, the concept that omental adipocytes facilitate OC spread in patients has been discussed^19^. Relatedly, a possible link between obesity-induced metabolic syndrome and OC has been made^20,21^. Because it is difficult to directly study adipocyte content in vivo clinically, numerous translational studies have examined body mass index and other correlates of visceral adiposity instead. The use of these metrics is limited when OC patients present with ascites, tumor masses, or small bowel obstructions that can affect absolute body weight and mask underlying alterations in body composition^22^. While some studies have shown that obesity may influence chemotherapeutic potency^23–25^, most recent clinical data indicate that, ultimately, adiposity is not strongly associated with OC risk, prognosis, or survival^22,26–32^.

Deficiency of FABP4 in mice is not equivalent to loss of mature adipocytes. Therefore, the concept that adipocytes drive ovarian tumor cell progression in the peritoneal cavity remains untested. In the present study, using a mouse model devoid of mature adipocytes in the peritoneal cavity, we directly explored whether mature adipocytes are pivotal in the ability of OC to seed and grow in the omental or peritoneal compartment after tumor cells are placed in the peritoneal cavity to model how OC takes hold after metastasis to the peritoneal cavity. The absence of mature adipocytes did not impact omental tumor growth in the short or long term, nor did it influence therapeutic efficacy. Instead, our data suggest that endothelial cells within the omental vasculature are the primary sources of the fatty acid transport protein FABP4.

## Methods

### Mice

Adiponectin^Cre^ mice (The Jackson Laboratory, Bar Harbor, ME, USA: 028020, kindly provided by Dr. Steven Teitelbaum) were crossed with lox-stop-lox-Rosa26 diphtheria toxin (DTA) mice (The Jackson Laboratory: 010527: RRID: IMSR_JAX:028020), resulting in the ablation of mature adipocytes and consequent lipoatrophy. These lipoatrophic mice, which constitutively lack brown and white adipose tissue, and their littermate controls were maintained at thermoneutrality (30°C) from birth on a 12-h light/dark cycle. CD19^Cre^ mice (Jackson Laboratory, Bar Harbor, ME, USA: 006785: RRID: IMSR_JAX:006785) were crossed with Rosa-TDTomato^fl/fl^ mice (Jackson Laboratory, Bar Harbor, ME, USA: 007914: RRID: IMSR_JAX:007914) to generate CD19^Cre/+^TD reporter mice, in which cells derived from B cell lineage were Tomato^+^. Triple knockout B6.129S-Rag2^tm1Fwa^, Cd47^tm1Fpl^, Il2rg^tm1Wjl^/J mice (The Jackson Laboratory, Bar Harbor, ME, USA: 025730: RRID: IMSR_JAX:025730) and age-matched C57BL/6J (The Jackson Laboratory, Bar Harbor, ME, USA: 000664: RRID: IMSR_JAX:000664) were purchased directly. Tie2-Cre^+^ Fabp4^fl/fl^ mice and Lyz2-Cre+ Fabp4^fl/fl^ mice were provided through MTA with Gökhan S. Hotamişligil, who generated these mice^33^. Animals were provided a standard chow diet (LabDiet 5053 – PicoLab Rodent Diet 20) and water ad libitum. All experiments and procedures were conducted in accordance with procedures and protocols approved by the Washington University in St. Louis Institutional Animal Care and Use Committee.

### Fat Transplantation

Mature gonadal, inguinal, and axillary adipose depots collected from donor C57BL/6J WT female mice or littermates, approximately 14-18 weeks old, were transplanted into 7-10 week old lipoatrophic Adiponectin^Cre/+^ RosaDTA^fl/+^ female recipient mice, termed **d**istal **a**dipocyte **r**eceiving **l**ipoatrophic (DARL) mice post-surgery. The corresponding littermate control mice born with standard fat depots received parallel sham surgeries and are referred to as **p**eritoneal **a**dipocyte-**r**eplete **c**ontrol (PARC) mice post-sham surgery. The surgeries were conducted as previously described^34,35^ with a few modifications. After collection, the fat pads were placed in sterile saline on ice until the recipient mouse was prepared for surgery. The lipoatrophic mice and littermate controls were anesthetized by continuous inhalation of 1-2% isoflurane and placed on a heating pad in the prone position. A small 1-2 mm incision was made in the dorsal scruff of transplant-receiving mice. Three small pockets were made bilaterally. In the lipoatrophic mice, one graft (150-300 mg) was implanted subcutaneously in each pocket, totaling six fat pads per mouse. In the littermate control mice, these six pockets were made but not engrafted. The incision was closed with 5-0 nylon sutures. The mice were given a subcutaneous injection of extended-release buprenorphine (1 mg/kg) and placed overnight in an infant incubator warmed to 27°C. Then, all mice were housed individually for a week to facilitate recovery before returning to five mice per cage.

### Omentectomy

Mice were anesthetized by continuous inhalation of 1-2% isoflurane and placed on a heating pad in the supine position. A small incision was made midline in the skin and then through the linea alba, a thin midline band of connective tissue along the peritoneal wall, to avoid bleeding. The greater omentum, including its associated mesothelial sheet, was carefully exposed and cauterized through this incision. The blood vessels supplying the omentum were cauterized to avoid bleeding. The peritoneal incision was closed with 4-0 Vicryl absorbable continuous sutures. The skin was closed with non-absorbable 5-0 nylon sutures. The mice were given a subcutaneous injection of extended-release buprenorphine (1 mg/kg) and placed in an infant incubator warmed to 27°C overnight for recovery before returning to room temperature.

### Tumor Cell Culture

The original ID8 parental cell line, derived from C57BL/6 mouse (RRID: IMSR_JAX:000664) ovarian surface epithelial cells and transfected with firefly luciferase, was directly shared via MTA with Katherine Roby, who developed the cell line^36^. These ID8 tumor cells labeled with firefly luciferase (ID8-Luc) recapitulate high-grade serous ovarian cancer (HGSOC) dissemination upon intraperitoneal injection in immunocompetent mice. Patients with HGSOC frequently have germline mutations in BRCA and p53 genes^37^. Hence, ID8p53^−/−^Brca2^−/−^ cells labeled with green fluorescent protein (GFP) and firefly Luciferase (Luc) were kindly donated by the Khabele lab (WashU) through an MTA with the McNeish lab^38^. The ID8p53^−/−^Brca2^−/−^ GFP Luc cells were maintained in high-glucose DMEM supplemented with 4% FBS, 1% Insulin-Transferrin-Selenium X, and 1% penicillin/streptomycin. The BPPNM (*Trp53*^−/−*R172H*^*Brca1*^−/−^*Pten*^−/−^*Nf1*^−/−^*Myc*^OE^ genotype) and KPCA (*Trp53*^−/−R172H^*Ccne1*^OE^*Akt2*^OE^*KRAS*^G12V^) murine fallopian tube epithelial cells^39^ were obtained by MTA with Robert Weinberg. The cells were maintained in DMEM containing 4% FBS, 2 ng/mL epidermal growth factor, 1% Insulin-Transferrin-Selenium X, and 1% penicillin/streptomycin. Modified BPPNM-luc cells and sorted KPCA-luc cells, which express firefly Luciferase homogenously, were provided by MTA with Nan Zhang. All cells were sub-cultured twice before each tumor injection. Cells were tested every six months for mycoplasma.

### Tumor Expansion In Vivo

Mice were injected intraperitoneally (i.p.) with 5×10^6^ ID8-Luc, 5×10^6^ ID8p53^−/−^Brca2^−/−^ GFP Luc, 1×10^6^ KPCA, or 3×10^6^ BPPNM tumor cells in 200 µl of sterile Hanks’ Balanced Salt Solution (HBSS) to mimic HGSOC dissemination in the peritoneal cavity. Bioluminescence imaging was performed weekly to quantify tumor burden in the peritoneal cavity noninvasively. Mouse weights were tracked weekly, and mice were sacrificed within a week of the appearance of detectable ascites at the latest. Mice were treated weekly with i.p. 30 mg/kg carboplatin (Selleckchem, Cat#S1215) as indicated.

### In Vivo Bioluminescence Imaging

In vivo bioluminescence imaging was performed on the days indicated using the IVIS50 system (PerkinElmer, Waltham, MA) with Living Image 4.3.1 up to 1min exposure(s), eight bin, FOV12cm, f/stop1, and open filter. Mice were injected i.p. with D-luciferin (150mg/kg in PBS; Gold Biotechnology, St. Louis, MO) based on mouse weight and imaged 10 minutes later while under isoflurane anesthesia (2% vaporized in O_2_). Total photon flux (photons/sec) was measured from fixed regions of interest (ROIs) over the peritoneal cavity using Living Image 2.6 (RRID: SCR_014247).

### Ex Vivo Bioluminescence and Stereoscope Imaging for Tumor Quantification

Immediately before sacrifice, tumor-bearing mice were injected i.p. with D-Luciferin (150mg/kg in PBS; Gold Biotechnology, St. Louis, MO) based upon their weight. Mice were sacrificed 10 minutes post-injection. Peritoneal organs were washed in PBS and pinned in Sylgard plates containing CO_2_-independent media with 5% FBS on ice. When the plates were ready to be imaged on a ChemiDoc MP Imaging System (Bio-Rad) or the IVIS50, the media was replaced with fresh CO_2_-independent media containing diluted D-Luciferin (1:100) to ensure an equivalent duration in the substrate across all samples and maintain a stable pH. The plates were warmed at room temperature until the temperature equilibrated for optimal signal detection^40^.

Images generated from the IVIS50 and the corresponding Living Image software were used to quantify the tumor burden in each organ. Because each organ was different in size, unique regions of interest (ROIs) were created that matched the size of each organ. The total photon flux (photons/sec) was normalized to the specific area of that organ, resulting in photon flux/area measurements.

Chemiluminescence and colorimetric images generated in the ChemiDoc reader were used to quantify the number of tumor nodules in each organ’s adipose or sheet region. Images were imported into FIJI software (RRID: SCR_003070) for analysis. A global scale was set across all the images to quantify the nodules’ size in cm. The auto-threshold features binarized the chemiluminescence images to distinguish the nodules from the background, thereby eliminating user bias. The analyze particles feature was used to determine the number of nodules, the size of each nodule, and the total area covered by nodules. The counted particles or nodules were then superimposed on brightfield stereoscope images to determine whether they were localized to each tissue’s adipose or sheet regions, allowing us to quantify the percentage of the fat or sheet region occupied by tumor nodules for each organ.

### Two-photon Microscopy

CD19^Cre/+^TD reporter mice injected with ID8p53^−/−^Brca2^−/−^ GFP Luc cells were anesthetized with ketamine/xylazine cocktail (0.1 mL/25 g). The omentum was exposed through a 1-2mm incision in the skin and peritoneum. The mice or fixed omentum were positioned under the customized Leica SP8 2-Photon microscope equipped with a 325/0.95 NA water-dipping objective and a Mai Tai HP DeepSee Laser (Spectra-Physics) tuned to 810 nm. Fluorescence emission was separated by 3 high-efficiency dichroic mirrors cutting at 458, 495, and 560 nm (Semrock) and directly directed to 4 supersensitive external detectors.

### H&E Staining

Tissue samples were fixed overnight in 4% paraformaldehyde and transferred to PBS the next day. The samples were processed and stained with hematoxylin and eosin by iHisto (Boston, Massachusetts).

### Serum Triglyceride Measurements

At 5-6 weeks post adipose tissue transplant, serum was collected from whole blood via cheek bleed in BD Microtainer® Blood Collection Tubes with Serum Separator Gel. The serum was allowed to clot at room temperature for approximately 60 minutes. The clot was removed by centrifugation at 1500 RCF for 10 minutes at 4°C. Following centrifugation, the serum was collected. Serum triglycerides were measured per the manufacturer’s instructions (Wako L-Type TG M), and the remaining samples were stored at -80°C.

### Glucose and Insulin Tolerance Tests

Serial blood glucose measurements were taken from venous blood (Glucocard, Vital) at 0, 15, 30, 60, and 90 minutes following intraperitoneal injection of D-glucose (1.5g/kg) or insulin (0.75 U/kg). Mice were fasted for six hours prior to the insulin tolerance test or overnight for the glucose tolerance test. Mice were weighed before and post-fasting. During fasting, mice were placed in clean cages with a metal grid on the bottom to prevent them from making contact with the cage bottom, thereby limiting coprophagy. To calculate the area of the curve (AOC), the glucose measurements were plotted, and the area under the baseline was subtracted from the area under the curve.^41^

### FABP4 ELISA

Mouse FABBP4 was measured in plasma using a commercially available ELISA kit (Abcam, Cat# ab277426).

### In Vivo Antibody Depletion

Anti-Mouse CD4 (Leinco Technologies Cat# C1333, RRID: AB_2737452), anti-Mouse CD8b.2 (Leinco Technologies Cat# C2832, RRID: AB_2737471) and anti-Mouse NK1.1 (Leinco Technologies Cat# N123, RRID: AB_2737553) purified functional-grade *in vivo* gold antibodies were purchased from Leinco. Starting one week before cancer injection, mice were intraperitoneally injected weekly with 250 µg of the CD4 (RRID: AB_2737452) and CD8 (RRID: AB_2737471) depleting antibodies and 100 µg of the NK depleting (RRID: AB_2737553) antibody. Antibody depletion was validated throughout the experiment using flow cytometry of plasma blood samples collected via cheek bleeds.

### Murine Mesentery and Omentum Tissue Collection and Sheet Separation

Whole mesenteries and omenta from PARC and DARL mice, eight weeks post-surgery, were pinned onto Sylgard dishes and kept hydrated in PBS. The adipose region in PARC mice and the corresponding area in DARL mice were carefully separated from the membranous sheet region for each omentum and mesentery using a scalpel under a dissection microscope and placed in RPMI with 5% FBS on ice. Each sheet or adipose region collected from every mouse was minced with scissors and digested separately using 1 mg/mL collagenase IV, 0.1 mg/mL DNAse, and 0.5% BSA for 30 minutes on a shaker at 150 rpm. The samples were centrifuged and resuspended for downstream processing.

### Human Omentum Tissue Collection and Processing for Single Cell Sequencing

Benign omentum was collected under IRB approval number 201903051 via the Washington University Ovarian Cancer Tissue Bank. Using a scalpel, the tissue was dissected into omentum sheet and fat regions, which were then minced separately and transferred into tubes for enzymatic digestion. Digestion media consisted of RPMI 1640 supplemented with 1 mg/mL collagenase IV and 0.1 mg/mL DNase, and samples were incubated for 45 minutes at 150 rpm. Digestion was then stopped by adding chilled FACS buffer, and the cell suspension was passed through a 70 µm filter before centrifugation at 500 g for 5 minutes at 4°C. Cell pellets were resuspended in FACS buffer containing DAPI, and DAPI-negative (live) cells were sorted. All sorted live cells were then submitted for single-cell RNA sequencing. RNA libraries were prepared using the Chromium Next GEM Single Cell 5’ Kit v2 (10x Genomics, #1000263) according to the manufacturer’s instructions. In brief, cells were encapsulated into droplets, barcoded during reverse transcription, and the emulsions were broken to enable cDNA amplification via PCR, following the V(D)J enrichment protocol (10x Genomics, #1000252). Libraries were prepared according to 10x Genomics guidelines and sequenced on an Illumina NovaSeq 6000 platform, with samples pooled and barcoded to ensure sufficient read depth. Raw sequencing data were processed using Cell Ranger (versions 3.1.0, 6.1.1, or 7.0.1) with the GRCh38 human reference genome to produce gene expression matrices from single-cell 5′ RNA-seq data.

### Spectral Flow Cytometry

Mesentery and omentum samples were prepared as described (see Mesentery and Omentum Tissue Collection and Sheet Separation). Blood samples were collected in tubes containing 15ml of 500mM EDTA, and red blood cells were lysed using BD PharmLyse lysis buffer. Blood cells were stained with acridine orange, and leukocytes were counted on a Nexcellom Cellometer Auto X4 cell counter. Quantifying omental and mesothelium-associated cells was based on cytometer counts provided by the Cytek Aurora spectral cytometer. For flow cytometry analysis, cells were washed in PBS, stained with ZombieNIR (cat# 423106), washed again, and resuspended with the antibody mix diluted in flow cytometry buffer (HBSS containing 2% FBS and 5mM EDTA) containing 20% BD Horizon^TM^ Brilliant Stain Buffer (cat#563794) for 30 min at 4°C. The flow antibodies used at a concentration of 1:300 were as follows: CD45 (BD Biosciences Cat# 748370, RRID:AB_2872789), F4/80 (BioLegend Cat# 123168, RRID: AB_2924461), CD64 (BioLegend Cat# 139313, RRID: AB_2563903), ICAM-2 (BD Biosciences Cat# 740864, RRID:AB_2740516), CD169 (BioLegend Cat# 142416, RRID: AB_2565620), Lyve1 (Thermo Fisher Scientific Cat# 53-0443-82, RRID: AB_1633415), MHCII (BD Biosciences Cat# 750281, RRID: AB_2874472), CD31 (BD Biosciences Cat# 741262, RRID: AB_2870809), CD11b (BD Biosciences Cat# 564443, RRID: AB_2738811), Ly6g (BioLegend Cat# 127641, RRID: AB_2565881), Ly6c (BioLegend Cat# 128029, RRID: AB_10896061), Cd11c (Thermo Fisher Scientific Cat# 363-0114-82, RRID: AB_2925254), Nk-1.1 (BioLegend Cat# 156519, RRID: AB_2894654), CD4 (BioLegend Cat# 100467, RRID: AB_2734150), CD8a (BioLegend Cat# 100791, RRID: AB_2876397), CD19 (clone 1D3), catalog # 50-245-964), podoplanin (BioLegend Cat# 127411, RRID: AB_10613294), PDGFR-α or CD140a (BioLegend Cat# 135913, RRID: AB_2715986), and CD9 (BD Biosciences Cat# 751253, RRID: AB_2875270). Cells were washed twice in flow bluffer before data acquisition. Flow cytometry data were acquired on a Cytek Aurora spectral cytometer (5 laser configuration). Flow cytometry data analysis, including unsupervised analysis, was conducted using FlowJo software v10.10 (RRID:SCR_008520).

### Cell Hashing and Sorting for Murine Single-Cell RNA Sequencing

Following digestion of the omentum and mesentery from PARC and DARL mice, the samples were washed and passed through 100 μm filters and stained with CD45 (AB_312980, BioLegend Cat. No. 103115), DAPI, and TER-119 (AB_313706, BioLegend Cat. No. 116205) antibodies in one antibody mixture for all samples. Each sample was then individually stained with one of 8 hashtags or antibody-derived tags (ADTs). The eight TotalSeq antibodies used were AB_2800693 (BioLegend Cat. No. 155861), AB_2800694 (BioLegend Cat. No. 155863), AB_2800695 (BioLegend Cat. No. 155865), AB_2800696 (BioLegend Cat. No. 155867), AB_2800697 (BioLegend Cat. No. 155869), AB_2819910 (BioLegend Cat. No. 155871), AB_2819911 (BioLegend Cat. No. 155873), and AB_2819912 (BioLegend Cat. No. 155875). These ADTs are conjugated with a DNA oligo containing a barcode to target distinct ubiquitous cellular surface epitopes and were used to identify independent biological replicates for each tissue. The samples were washed three times after hashtag staining to remove unbound antibodies before being pooled into four separate tissue groups, mesenteric fat region (MFR), mesenteric sheet (MS), omentum fat region (OFR), and omentum sheet (OS), for both PARC and DARL mice before sorting. Samples were kept on ice until sorted on the FACSAria II with a 100-μm nozzle (Becton Dickinson). Cells positively stained with DAPI were gated out during cell sorting to remove dead cells. Similarly, cells positively stained for TER-119 were excluded to remove red blood cells from our cell quantifications and increase the collection of desired cells. Approximately equal quantities of CD45^+^ and CD45^-^ cells were submitted for 10X Genomics single-cell RNA sequencing for each sample via the McDonald Genome Institute Core.

### Murine Single-Cell RNA Sequencing

Raw gene expression matrices were processed using the Seurat package in R (V5.1.0). Cells expressing at least 200 genes and genes present in at least three cells were included in the analysis. Antibody capture data from the ADTs were added as a separate assay to facilitate multiplexing analysis, and cell identifiers were renamed to ensure uniqueness across all samples. Single-cell BCR and TCR data were also acquired. The individual datasets were merged into a single Seurat object for integrated analysis. Following sequence alignment, counts of these tags were used to assign a hashtag to each cell. The standard Seurat workflow developed by the original authors of this technology was used to demultiplex hashtag oligos (https://satijalab.org/seurat/articles/hashing_vignette.html). Cells with no hashtags (negatives) or multiple hashtags (multiplets) were removed from the analysis. Quality control metrics were calculated, and cells with more than 50,000 counts, fewer than 200 features, more than 6,000 features, or mitochondrial gene expression exceeding 15% were excluded. Data normalization was performed, and 3,000 highly variable features were identified using the variance-stabilizing transformation method. To correct for batch effects, the Harmony integration algorithm was applied using principal component analysis for dimensionality reduction. The top 30 principal components were used for downstream analysis. Clustering was performed over a range of resolutions to identify cell populations at various levels of granularity. Dimensionality reduction for visualization was achieved using Uniform Manifold Approximation and Projection (UMAP) on the integrated data. Metadata was updated to include genotype (Cre-negative or Cre-positive), tissue type (omentum or mesentery), and region (sheet or fat) based on the sample origin. UMAP plots were generated to visualize the distribution of cells grouped by these categories. The fully processed and integrated dataset was saved for subsequent analyses.

### Capybara Analysis and Custom Reference Generation

Capybara analysis was performed to evaluate relative cell identity with several custom references (R package version 0.0.0.9; https://github.com/morris-lab/Capybara). Tissue-level classification was not executed; instead, custom references were generated using the Capybara function *construct.high.res.reference()*. Published datasets were selected as appropriate references for this dataset for a couple of reasons: they contain cell types and tissues in common with our data, identified by rigorous cell type classification by the original authors of each dataset, and they contain a variety of cell types that are expected to represent the diversity of cell types in our dataset as fully as possible. Following reference construction, quadratic programming, discrete cell identity classification, and multiple identity scoring were performed on our query dataset with each reference independently.

Two publications made both the counts and original cell type classifications publicly available both for their single-cell data and analysis ^42,43^. Raw counts and cell metadata for Brulois *et al.*^42^ were obtained from GEO (GSE140348). Raw counts and cell metadata are available for Merrick *et al*.^44^ and Rivera-Gonzalez *et al*. on FigShare (RRID: SCR_004328) (https://tinyurl.com/5et2bh5n); raw counts for these datasets are available on GEO (GSE128889 and GSE 241627, respectively). Where necessary, cell type classification efforts were recapitulated according to the original criteria and methods (See supplemental methods of Rivera-Gonzalez *et al*.^43^ for cell type classification of Merrick *et al.* ^44^).

### Human Single-Cell RNA Sequencing

Raw gene expression matrices were processed using the Seurat package in R (V5.1.0). Count matrices from each library were read in with Read10X and converted to Seurat objects using CreateSeuratObject (min.cells = 3, min.features = 200). Cells were renamed to ensure unique barcodes across samples by removing trailing “-1” suffixes, and individual Seurat objects were merged into a single object for joint analysis. The percentage of mitochondrial gene expression was calculated using PercentageFeatureSet(pattern = “^MT-”), and cells were filtered based on quality control metrics: only cells with fewer than 25,000 total counts, more than 500 and fewer than 6,000 detected features, and less than 10% mitochondrial reads were retained. Filtered data were normalized with NormalizeData, and the top 3,000 highly variable genes were identified using FindVariableFeatures. Data were scaled with ScaleData while regressing out nCount_RNA and percent.mt, and principal component analysis was performed with RunPCA. The first 25 principal components—accounting for approximately 88% of variance—were used to construct a shared-nearest-neighbor graph (FindNeighbors, dims = 1:25) and to perform graph-based clustering across resolutions 0.1 to 1.5 (FindClusters). UMAP embeddings were computed on these 25 PCs (RunUMAP, dims = 1:25) to visualize cell populations. Cluster identities and original sample labels (orig.ident) were preserved in metadata for downstream interpretation. UMAP plots were generated for each clustering resolution as well as grouped by orig.ident, and the fully processed Seurat object was saved for subsequent analyses.

### Statistics

The number of mice used in each experiment and the statistical analyses performed are indicated in the figure legends. Smaller pilot experiments informed the technical feasibility of experiments and the number of mice necessary to achieve sufficient statistical power. Co-housed age-matched littermate controls were used for all in vivo murine experiments with genetically altered mice. All littermates and genetically altered mice were treated identically and simultaneously during all experimental steps to remove confounding variables. Mice in each genotype were randomly divided into treatment or control groups when necessary. Parametric statistical tests were used for statistical analysis. Thus, distributions were assumed to be normal, but this was not formally tested. Error bars represent standard error. Statistical significance was evaluated using two-tailed unpaired Student’s *t*-tests or two-way ANOVAs followed by Tukey’s Honestly Significant Difference (HSD) Test for pairwise multiple comparisons. A Chi-Square Test was performed on the capybara distribution analyses for each reference to determine if the cell type distribution varied between genotypes (α= 0.05). Differential expression analysis was performed using the default parameters of the Seurat function *FindMarkers()*. These include requirements of a 0.25 log fold-change between the two groups, a minimum fraction of .1 cells expressing the feature in either population for testing, and the use of a Wilcoxon Rank Sum test to identify features. All statistical calculations were performed with GraphPad Prism 10 (RRID: SCR_002798). A *p-value* < 0.05 was considered statistically significant.

## Results

### Ovarian cancer spatially seeds regions of adiposity within peritoneal organs

After intraperitoneal (i.p.) injection of ID8p53^−/−^Brca2^−/−^ GFP Luc tumor cells, comprehensive biodistribution studies were conducted at 7 days (Extended Data Fig. 1a) and six weeks (Extended Data Fig. 1b) to comprehensively pinpoint the organs that tumor cells consistently seed for further analysis. We compared tumor burden in the omentum to that in other organs, including other organs endowed with visceral white adipocytes in the peritoneal cavity, such as the mesentery, perirenal adipose depot, and gonadal fat pads. Fat pads outside the peritoneal cavity, such as the axillary and inguinal fat pads, and organs in the thoracic cavity, including the heart and lungs, showed little to no signal, indicating that the tumor burden was predominantly restricted to the peritoneal cavity, as expected, given the route of tumor administration. The pancreas, located near the omentum, increased in tumor burden over time. Other peritoneal organs, such as the stomach, spleen, small intestines, colon, cecum, kidneys, adrenal gland, and liver, maintained low signal throughout the tumor course (Extended Data Fig. 1a, b). Overall, the five most common sites of metastasis consistently observed were the omentum, small bowel mesentery, peritoneal sheath, gonadal fat pads, and the diaphragm that separates the peritoneal and thoracic cavities. The omentum consistently had 2-3 times greater tumor burden than any other location, both early (Fig. 1a) and late (Fig. 1b), after tumor seeding. This pattern fits with clinical results reported in OC patients^2^. These five locations were consistently analyzed across all future experiments.

**Figure 1.**
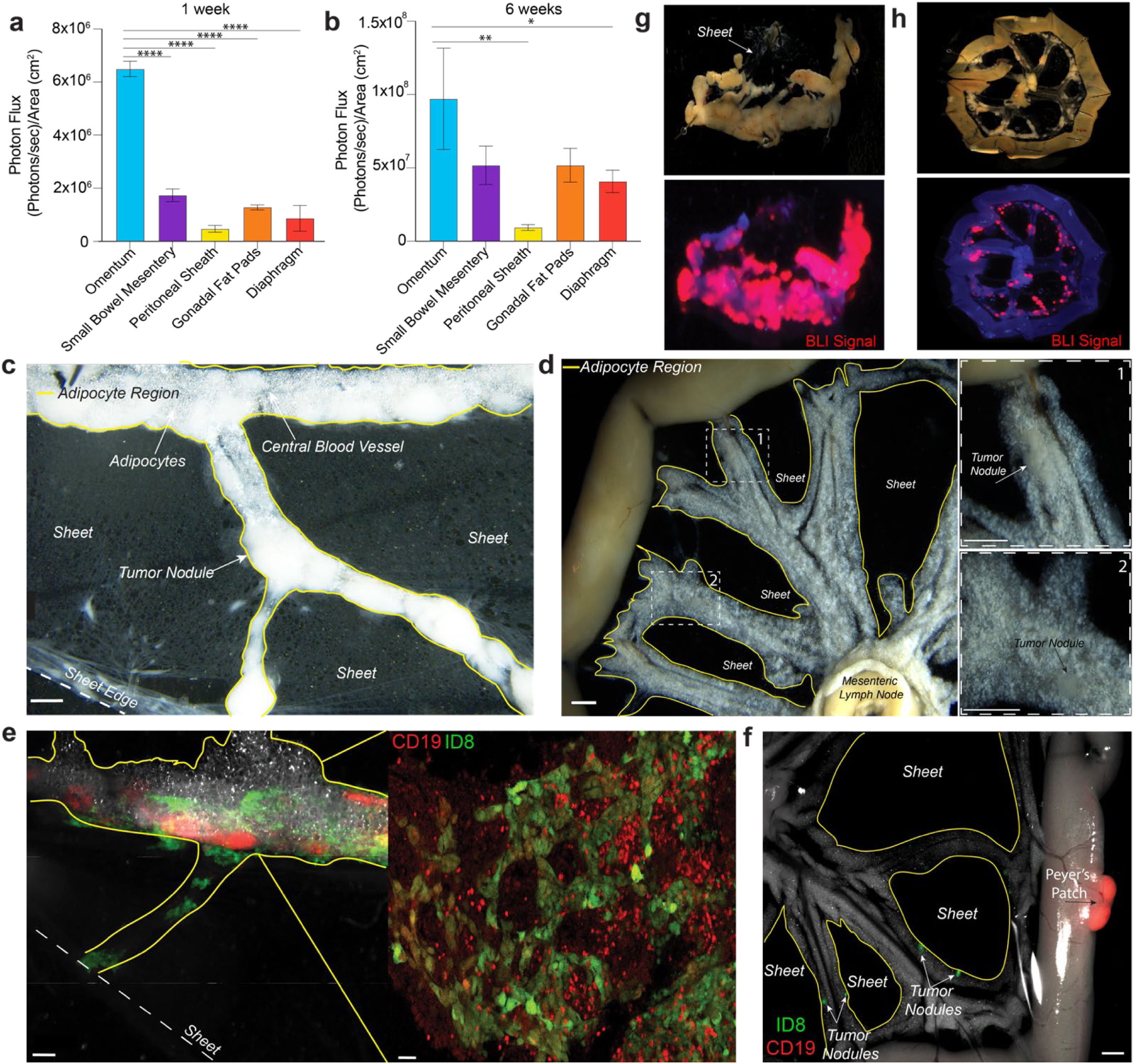
OC cells seed regions of adiposity in peritoneal organs. **a.** One week (*n*=5, One-way ANOVA****P<0.0001) and **b**. Six weeks (*n*=5, One-way ANOVA *P<0.05) post i.p. injection of 5×10^6^ ID8p53^−/−^Brca2^−/−^ GFP Luc tumor cells in C57BL/6J wild-type (WT) mice. Tukey’s HSD Test, ****P<0.0001, **P<0.01, *P<0.05. **c.** Representative brightfield stereoscope image of an omentum one week post i.p. injection of 5×10^6^ ID8p53^−/−^Brca2^−/−^ GFP Luc tumor cells. The adipocyte region is outlined in yellow. Scale bar, 1mm. **d**. Representative brightfield stereoscope image of a mesentery one week post i.p. injection of 5×10^6^ ID8p53^−/−^Brca2^−/−^ GFP Luc tumor cells in WT mouse. The adipocyte region is outlined in yellow. Scale bar,1mm. **e**. Stereoscope (left panel, scale bar, 200 μm) and focused 2-photon image (right panel, scale bar, 40 μm) of ID8p53^−/−^Brca2^−/−^ GFP Luc tumor cells in the adipocyte region of the omentum in a CD19^Cre/+^TD mouse three days post i.p. tumor cell injection**. f.** Stereoscope image of tumor nodules in the mesentery of a CD19^Cre/+^TD mouse three days post i.p. injection of 5×10^6^ ID8p53^−/−^Brca2^−/−^ GFP Luc tumor cells. Scale bar, 1mm. Corresponding brightfield stereoscope image (top) and ChemiDoc MP Imaging System bioluminescence (BLI) image (bottom) of the **g**. Omentum and h. Mesentery six weeks post i.p. injection of 5×10^6^ ID8p53^−/−^Brca2^−/−^ GFP Luc tumor cells in WT mice. BLI signal corresponding to tumor burden is shown in red, and blue is used for post-hoc contrast.

Having established that tumor burden had a propensity for peritoneal tissues containing adipocytes, we examined tumor seeding within these adipocyte-rich compartments more closely. The omentum is covered by a continuous mesothelial layer, and the underlying tissue can be anatomically partitioned into two regions: an adipose-rich, vascularized compartment and a translucent, membranous sheet largely devoid of adipocytes but composed of fibroblasts, mesothelial cells, and macrophages^45^. These distinct tissue types are macroscopically and histologically identifiable in both humans and mice^3^. In the present study, we leverage this anatomical framework to map tumor localization within the omentum and to examine how regional structure may influence metastatic seeding. By aligning our analysis with known anatomical features, we provide a detailed and functionally relevant characterization of the regional omental structure in the context of OC tropism.

After acute tumor seeding, these adipose and sheet regions in the mouse omentum are demarcated in Fig 1c. Similar variations between the adipocyte and nonadipocyte sheet regions characterize the human omentum^3,46^. Like the omentum, the mesentery is also organized into regions enriched in adipose tissue or translucent cellular tissue sheets (Fig. 1d) with a similar composition of fibroblasts, macrophages, and mesothelial lining cells^8,47^. While the omentum is associated with well-developed lymphoid aggregates called milky spots, the mesentery also contains fat-associated lymphoid clusters that increase in number and size with age^48^ and inflammation^49^. Thus, given their structural similarities, mutual location in the peritoneal cavity, and susceptibility to tumor seeding, we reasoned that the mesentery served as an ideal comparator for the omentum to explore preferential omental tumor homing within the peritoneal cavity.

In just days after i.p. injection of ID8p53^−/−^Brca2^−/−^ GFP Luc tumor cells in CD19^Cre/+^Tomato (TD) mice, stereomicroscopy (Fig. 1e, left) or intravital two-photon imaging (Fig. 1e, right) demonstrated that the GFP^+^ ID8 tumor cells had already migrated to the adipocyte region in the omentum, where CD19^+^ B cells demarcated the lymphoid-rich milky spots (Fig. 1e, left), and formed lattices over adipocytes (Fig. 1e, right). Tumor cells localized near milky spots in the omental adipose tissue, while some fluorescent tumor cells specifically seeded the adipocyte regions of the mesentery at a distance from CD19+ Peyer’s Patches (Fig. 1f). GFP^+^ tumor nodules in the mesentery were mainly located where the branching vasculature was also situated in the mesenteric adipose (Fig. 1f). While the adipose tissue of both the omentum and the mesentery provided a niche for tumor cells, there was more tumor burden in the omentum than in the mesentery (Fig. 1a-h). Six weeks after tumor implantation, tumor nodules persisted and expanded in regions of adiposity in the omentum and the mesentery, as shown in the brightfield and corresponding ex vivo bioluminescence images (Fig. 1g, h). Overall, these data indicate that the adipose tissue region of the omentum, and to a lesser extent, the adipose tissue region of the mesentery, supports ovarian tumor seeding and growth.

### Mature adipocytes are not required for OC seeding and growth within the peritoneal cavity

To formally test the role of mature adipocytes in tumor seeding and expansion, we crossed mice expressing Cre driven by the adiponectin promoter with mice bearing lox-stop-lox-Rosa26 diphtheria toxin (DTA) alleles to generate congenitally lipoatrophic mice that systemically and completely lacked mature brown and white adipocytes from birth, retaining only adipocytes in the bone marrow, as previously described^50^. Since breeders carried one allele encoding Cre downstream of the adiponectin promoter and one WT allele, half of the progeny from the cross yielded littermate controls with standard adipose tissue development. Brightfield images of the gonadal fat pad region, inguinal fat pad region, mesentery, and omentum in lipoatrophic and littermate control mice indicate that even lipoatrophic mice, without mature adipocytes, have tangible tissue in the canonical regions defined by fat depots (Extended Data Fig. 2a). It has been estimated that only 30% of cells in adipose tissue are mature adipocytes, and the lipoatrophic mice should still have a network of mesothelial cells, fibroblasts, endothelial cells, macrophages, stem cells, and pre-adipocytes that comprise the tissues in these regions^51^.

Lipoatrophy is associated with metabolic complications, including hepatic steatosis, dysregulated glucose tolerance, and high circulating triglyceride levels due to a lack of fat depots where excess triglycerides can be stored^52–55^. To normalize these metabolic parameters while retaining a complete loss of mature adipocytes in the omentum and other tissues within the peritoneal cavity, we performed subcutaneous fat pad transplants into the scruff of the lipoatrophic mice, leaving the peritoneal cavity devoid of any mature adipocytes, while supplying an adipocyte-rich tissue to the mice systemically. Sham surgeries were also performed in the littermate control mice, which retained their standard fat depots in the peritoneal cavity. After fat transplant rescue surgery, we termed the Adiponectin^Cre+^DTA^fl/+^ mice **d**istal **a**dipocyte**-r**eceiving **l**ipoatrophic mice (DARL) and the sham surgery-receiving littermate control mice **p**eritoneal **a**dipocyte-**r**eplete **c**ontrol mice (PARC). When recovered weeks later, the engrafted fat resembled the typical appearance of healthy adipose tissue on H&E staining (Extended Data Fig. 2b) and expanded to more than double the starting weight at the time of transplant (Extended Data Fig. 2c). The mice tolerated the surgery well, as there were no significant changes in body weight after surgery (Extended Data Fig. 2d). Serum triglycerides normalized after surgery and became statistically equivalent to that of fat-bearing control littermates (Extended Data Fig. 2e). Lipoatrophic mice initially had elevated fasting glucose levels compared to littermate controls, which became more pronounced with increased fasting duration (Extended Data Fig. 2f, g). The generation of DARL mice reduced fasting glucose levels so that they resembled PARC mice to a greater extent (Extended Data Fig. 2f, g). As previously reported^53^, lipoatrophic mice had dysregulated responses to insulin and glucose; these aberrations were largely corrected after surgery, as indicated by insulin and glucose tolerance tests (Extended Data Fig. 2h-k).

After confirming that metabolic parameters normalized in the lipoatrophic mice by six weeks after fat transplant, PARC and DARL mice were injected i.p. with ID8p53^−/−^Brca2^−/−^ GFP Luc tumor cells. Unexpectedly, weekly bioluminescence imaging demonstrated no difference in tumor burden at any point in the experiment (Fig. 2a), with the endpoint chosen to avoid the onset of profound ascites. Regional tumor burden quantified by photon flux revealed that the tumor distribution between PARC and DARL mice was similar across organs (Fig. 2b). This mirrored the pattern observed in unmanipulated C57BL/6J WT mice (Fig. 1a, b), with the omentum having the highest tumor burden, followed by the mesentery. Thus, the propensity of tumor cells to seed and expand in the omentum was prominent even in the absence of mature adipocytes.

**Figure 2.**
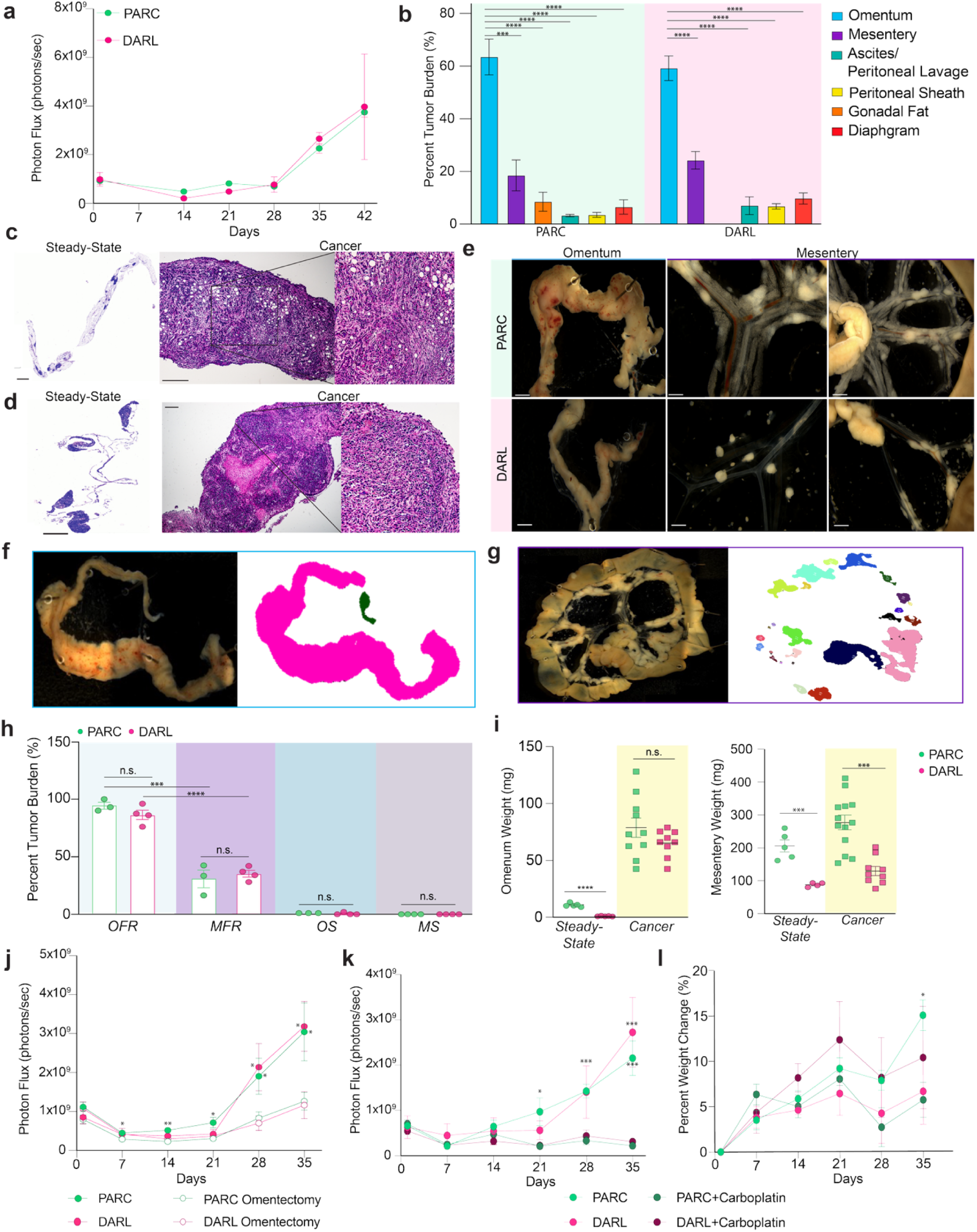
Omental adipocytes are not required for OC expansion in the peritoneal cavity. **a**. Quantification of in vivo bioluminescence images of PARC(*n*=4) and DARL(*n*=4) mice post i.p. injection of 5×10^6^ ID8p53^−/−^Brca2^−/−^ GFP Luc tumor cells. A two-way ANOVA (n.s. for genotype) and post-hoc comparisons using Tukey’s HSD were conducted to compare the means of the groups over time, n.s at any time point. **b**. Distribution of tumor burden across peritoneal organs based on quantifying ex vivo bioluminescence images of pinned organs in PARC (*n*=4) and DARL (*n*=4) mice six weeks after i.p. injection of 5×10^6^ ID8p53^−/−^Brca2^−/−^ GFP Luc tumor cells. Two-tailed Student t-test, ***P<0.001, ****P<0.001 **c**. H&E staining of steady-state PARC omentum (left, scale bar, 1 mm) and tumor-bearing omentum (six weeks post i.p. injection of 5×10^6^ ID8p53^−/−^Brca2^−/−^ GFP Luc tumor cells (right, scale bar 0.1 mm). **d**. H&E staining of steady-state DARL omentum (left, scale bar, 200 µm) and tumor-bearing omentum (six weeks post i.p. injection of 5×10^6^ ID8p53^−/−^ Brca2^−/−^ GFP Luc tumor cells (right, scale bar 0.1 mm). **e**. Stereoscope brightfield images of the omentum (first panel) and mesentery (second and third panels) in PARC and DARL mice six weeks post i.p. injection of 5×10^6^ ID8p53^−/−^Brca2^−/−^ GFP Luc tumor cells. Scale bars, 2 mm. **f**. Representative omentum brightfield image (left) and corresponding FIJI ImageJ tumor nodule masks (right) generated by particle analysis used to quantify the total area occupied by tumor nodules. **g**. Representative mesentery brightfield image (left) and corresponding FIJI ImageJ tumor nodule masks (right) generated by particle analysis used to quantify the total area occupied by tumor nodules, in which each tumor nodule is represented with a different color. **h**. Percent of the omentum fat region (OFR), mesentery fat region (MFR), omentum sheet (OS), and mesenteric sheet (MS) occupied by tumor in PARC (*n*=4) and DARL (*n*=4) mice six weeks post i.p. injection of 5×10^6^ ID8p53^−/−^Brca2^−/−^ GFP Luc tumor cells using measurements from FIJI Image J particle analysis. Two-Tailed Student’s t-test, ***P<0.001, ****P<0.0001. **i**. Weight of the omentum and mesentery from age-matched PARC and DARL mice at steady-state or six weeks post i.p. injection of 5×10^6^ ID8p53^−/−^Brca2^−/−^ GFP Luc tumor cells (cancer). Two-Tailed Student’s t-test ****P<0.0001, ***P<0.001. **j**. Quantification of in vivo bioluminescence images of PARC and DARL mice that received an omentectomy (PARC *n*=10, DARL *n*=8) or sham surgery (PARC *n*=9, DARL *n*=6). Mice were given six weeks to recover between their fat pad transplant and omentectomy surgeries (or corresponding sham surgeries) and six weeks post-surgery before i.p. injection of 5×10^6^ ID8p53^−/−^Brca2^−/−^ GFP Luc tumor cells. Two-Tailed Student t-tests between PARC and PARC omentectomy, **P<0.01, *P<0.05. Two-Tailed Student t-tests between DARL and DARL omentectomy, *P<0.05. **k**. Quantification of in vivo bioluminescence images of PARC and DARL mice treated with carboplatin (30 mg/kg i.p. weekly) (PARC, *n*=4, DARL, *n*=4) or saline (PARC, *n*=5, DARL, *n*=5). Two-way ANOVA followed by multiple comparisons with Tukey HSD between PARC and PARC treated with carboplatin or DARL and DARL treated with carboplatin, *P<0.05, ***P<0.001. **l**. Percent weight change of PARC and DARL mice treated with carboplatin (30 mg/kg i.p. weekly) after i.p. injection of 5×10^6^ ID8p53^−/−^Brca2^−/−^ GFP Luc tumor cells. Two-way ANOVA (n.s. for genotype) followed by multiple comparisons with Tukey HSD between PARC mice and PARC mice treated with carboplatin, *P<0.05.

We also evaluated other OC tumor models. We injected PARC and DARL mice i.p. with either 1×10^6^ KPCA-Luc cells or 3×10^6^ BPPNM-Luc cells (Extended Data Fig. 3), both derived from fallopian epithelial cells^39^. Representative endpoint images and tumor burden evaluation quantification of KPCA- and BPPNM-injected PARC and DARL mice (Extended Data Figure 3) showed that the omentum becomes encapsulated with tumor nodules in both genotypes, regardless of the presence of mature adipocytes (Extended Data Fig. 3a, c). The KPCA tumors quickly saturated the bioluminescence read-out and thus were harvested at day 7, when it was already clear that the omentum was a potent reservoir for tumor growth, regardless of the presence of mature adipocytes (Extended Data Fig. 3 a, b). BPPNM tumors, appearing to grow less aggressively than the KPCA model, were harvested at day 21, where again, the omentum was the dominant site of tumor residence regardless of the presence of mature adipocytes (Fig. 3c, d) These findings, consistent with results obtained using ID8p53^−/−^Brca2^−/−^ GFP Luc tumor cells (Fig. 2a, b), indicate that OC cells expand efficiently within the omentum in the absence of mature adipocytes, irrespective of tumor cell-of-origin (fallopian tube vs ovarian surface epithelium), genetic mutations, or the number of tumor cells injected.

**Figure 3.**
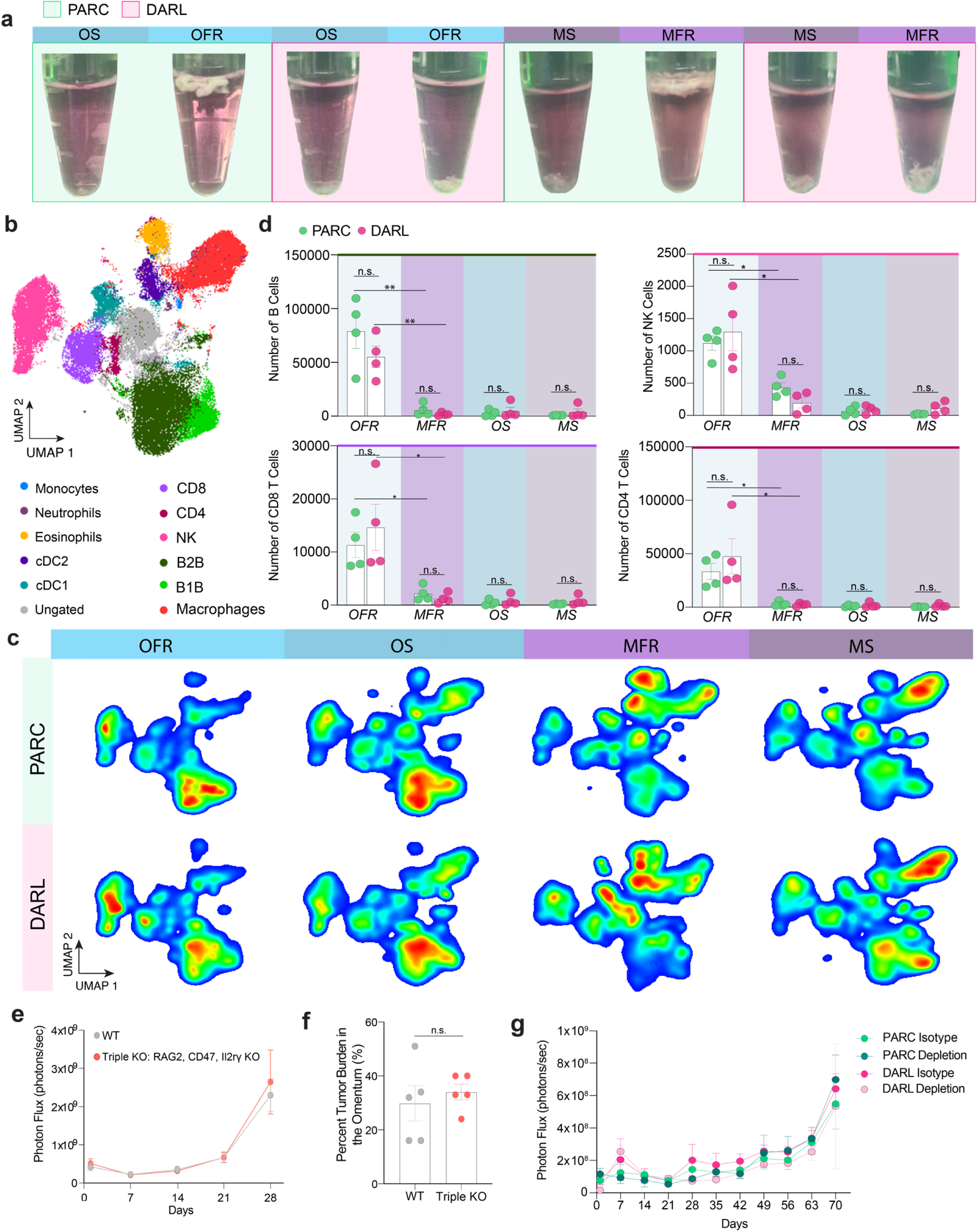
Immune characterization of lipoatrophic mouse model post metabolic normalization by fat transplant. **a**. Representative image following the separation of the sheet and fat regions in the omentum and mesentery of PARC and DARL mice to generate omentum fat region (OFR), omentum sheet (OS), mesenteric fat region (MFR), and mesenteric sheet (MS) distinct tissues. **b**. Dimensional reduction of CD45^+^ cells in PARC *(n*=4) and DARL (*n*=4) mice across all tissue samples (OFR, OS, MFR, MS) identified 12 distinct clusters based on spectral flow cytometry. **c.** Visualization of CD45^+^ population shifts in PARC (*n*=4) and DARL (*n*=4) mice across the OFR, OS, MFR, and MS. **d**. Quantification of the number of NK cells, CD8^+^ T cells, CD4^+^ T cells, and B cells in PARC (*n*=4) and DARL (*n*=4) mice across the OFR, OS, MFR, and MS. Two-Tailed Student’s t-test, *P<0.05, **P<0.01. **e**. Quantification of weekly in vivo bioluminescence images of age-matched WT (*n*=5) and Triple KO (*n*=5) mice purchased from The Jackson Laboratory after i.p. injection of 5×10^6^ ID8p53^−/−^Brca2^−/−^ GFP Luc tumor cells. One-way ANOVA (n.s. for genotype) followed by multiple comparisons with Tukey HSD, n.s. **f**. Percent tumor burden in the omentum of WT (*n*=5) and Triple KO (*n*=5) mice determined by quantification of ex vivo bioluminescence four weeks post i.p. injection of 5×10^6^ ID8p53^−/−^Brca2^−/−^ GFP Luc tumor cells. Two-Tailed Student’s t-test, n.s. **g**. Quantification of in vivo bioluminescence images of PARC and DARL mice treated weekly with CD8, CD4, and NK1.1 depleting antibodies (PARC depletion, *n*=5, DARL depletion, *n*=3) or isotype controls (PARC Isotype, *n*=5, DARL isotype, *n*=5) starting one week before i.p. injection of 5×10^6^ ID8-Luc tumor cells. One-way ANOVA (n.s.) followed by multiple comparisons with Tukey HSD, n.s.

Histological evaluation of the omentum from PARC mice after six weeks of ID8p53^−/−^Brca2^−/−^ GFP Luc tumor growth revealed few adipocytes remained by this time point, with tumor nodules surrounded by profound inflammation and fibrosis (Fig. 2c, right), while the DARL omentum completely lacked adipocytes (Fig. 2d). After pinning out the omentum and mesentery, it became apparent that the tumor nodules preferentially encapsulated the fat region of the omentum in both PARC and DARL mice (Fig. 2e) despite the latter lacking mature adipocytes. Similarly, tumor nodules formed along the adipose regions of the mesenteric branches, even when these regions lacked mature adipocytes in the DARL mice (Fig. 2e).

The tumor nodules in the adipose or sheet region of the omentum and mesentery were quantified using particle analysis tools in FIJI ImageJ software (Fig. 2f). In the PARC mice possessing mature adipocytes, the fat region of the omentum was supplanted by tumor nodules, defined by a few large tumor nodules encapsulating the entire structure (Fig. 2f). The fat region of the mesentery was also occupied by tumors but was comprised of numerous smaller nodules (Fig. 2g). In both DARL and PARC cohorts, nearly 100% of the omental fat region was occupied by tumor burden, even in the DARL mice lacking mature adipocytes (Fig. 2h). In contrast, less than 5% of the omentum sheet was colonized by tumors (Fig. 2h). Similarly, in both DARL and PARC cohorts, nearly 30% of the mesenteric fat region was occupied by tumors, while the sheet was virtually spared (Fig. 2h). Thus, while mature, lipid-laden adipocytes were not required to establish tumor growth, some other features of the fat region formed a niche particularly conducive to tumor seeding and proliferation. Due to the lack of mature adipocytes, the omentum and mesentery starting weights in the DARL mice were significantly decreased compared to their age-matched counterparts in the PARC mice at steady-state (Fig. 2i), which is also evident in the histological images of the omenta at steady state (Fig. 2 c, d, left). However, at the end of the experiment, once the tumors engrafted and expanded, the weights of the DARL omenta nearly matched the PARC omenta, indicating the equivalent dominance of the tissue by tumor nodules (Fig. 2i, left). In contrast, the weights of the DARL mesenteries were still substantially lower than the PARC mesenteries (Fig. 2i, right). These data indicate that the area of tissue normally housing lipid-laden adipocytes is highly hospitable to tumor growth. Still, the presence of lipid-laden adipocytes is dispensable for this effect.

We next wondered if the omentum was still critical for robust tumor expansion in the absence of mature adipocytes. To address this question, after a cohort of PARC and DARL mice recovered from sham or fat transplant subcutaneous surgeries, respectively, they were given omentectomies or sham peritoneal surgeries. Upon recovery from the omentectomies, the mice were injected i.p. with ID8p53^−/−^Brca2^−/−^ GFP Luc tumor cells. The omentectomized DARL and PARC mice had significantly reduced tumor burdens compared to those that received a sham peritoneal surgery, indicating that the omentum still plays a crucial role in metastasis even when it lacks mature adipocytes (Fig. 2j).

To consider the further possibility that adipocytes may impact therapeutic efficacy and tolerance, even if they are not required for tumor expansion, we treated PARC and DARL mice receiving ID8p53^−/−^Brca2^−/−^GFP Luc tumor cells with carboplatin, a chemotherapeutic agent that is part of the first-line standard of care for HGSOC^56^. This treatment resulted in an equivalent decrease in tumor burden between the PARC and the DARL mice (Fig. 2k), indicating that mature adipocytes do not influence the efficacy of chemotherapy. Additionally, the percent change in body weight between peritoneal tumor-bearing PARC and DARL mice treated with chemotherapy revealed no differences (Fig. 2i). Therefore, the lack of mature adipocytes does not alter chemotherapeutic efficacy or toxicity.

In conclusion, the absence of mature adipocytes had a minimal influence on tumor growth in the peritoneal cavity, regardless of whether omentectomy or chemotherapy was performed. Yet, the area where adipocytes usually reside remained the most conducive to tumor seeding and expansion. Even without mature adipocytes, the omentum remained a key driver of tumor expansion in the peritoneal cavity.

### Natural killer cell and lymphocyte deficiency do not impact omentum-centric OC growth in the peritoneal cavity

The omentum’s unique lymphoid aggregates, known as milky spots, play a role in adaptive immunity, potentially supporting the development of host defense against tumors or tumor-promoting Tregs^57^ that may influence tumor growth. We wondered if lacking mature adipocytes impacted the overall immune cell composition in the omentum or mesentery. Approximately ten weeks after surgery, the omentum and mesentery from eight DARL and PARC mice were collected. The tissues’ sheet and fat regions (Fig. 1c,d) were carefully separated before pooling tissue regions for spectral flow cytometry, generating four unique sample types: (i) omental fat region (OFR), (ii) mesenteric fat region (MFR), (iii) omental mesothelial sheet (OS), and (iv) mesenteric mesothelial sheet (MS). Samples containing mature adipocytes were readily identified because mature adipocytes floated (Fig. 3a), further confirming that the peritoneal cavity tissues from DARL mice lacked mature adipocytes.

Manual flow cytometry gating (Extended Data Fig. 4a) was used to identify CD45^+^ immune populations overlaid on a dimensionality reduction plot, identifying 12 clusters (Fig. 3b). Shifts in these clusters were visualized in OFR, OS, MFR, and MS dimensionality reduction plots (Fig. 3c). Among the CD45^+^ cells, natural killer (NK), CD4^+^, and CD8^+^ T cells appeared slightly elevated. In contrast, B1b cells were slightly reduced in the omentum fat region of DARL compared with PARC mice. Quantifications of these four cell types are shown in Fig. 3d. In contrast, the composition of the omentum sheet was similar across groups. Macrophages numbers with or without mature adipocytes did not obviously differ and were most abundantly retrieved in the mesenteric sheet (Fig. 3c). Mesenteric fat but not omental fat regions contained abundant eosinophils, many of which were still present in the DARL mice that lacked mature adipocytes (Fig. 3c).

To consider the possibility that the adaptive cytolytic immune compartment might influence ID8p53^−/−^ Brca2^−/−^ GFP Luc tumor cell seeding in the peritoneal cavity, we studied tumor growth in triple knockout mice that do not express genes encoding recombination activating gene 2 (Rag2), CD47, or the X-linked interleukin 2 receptor gamma chain (Il2rg) that comprises part of the receptor for several cytokines impacting lymphocyte and NK cell homeostasis. These mice, thus, genetically lack NK cells and T or B lymphocytes. When these mutant mice or control mice were i.p. injected with ID8p53^−/−^Brca2^−/−^ GFP Luc tumor cells, we observed no difference in the overall rate of tumor expansion between WT and triple KO mice (Fig. 2e). At the endpoint, importantly, there was also no difference in tumor burden in the omentum quantified by ex vivo BLI (Fig. 2f).

To reinforce this finding and to determine if the lack of adipocytes, coupled with the lack of T lymphocytes and NK cells, affected the tumor expansion, PARC and DARL mice were treated with depleting CD8, CD4, and NK1.1 antibodies after injection with the less aggressive ID8 parental tumor cell line labeled with firefly luciferase. Blood samples were analyzed via flow cytometry to confirm lymphocyte or NK cell depletion throughout the experiment (Extended Data Fig. 5). The tumor expansion itself depleted total NK cells but not T cells (Extended Data Fig. 5). These depleting antibodies used in PARC mice, replete with adipose tissue, recapitulated the lack of an impact on tumor expansion observed in the triple knockout mice. Conducting this immune cell depletion experiment in DARL mice lacking mature adipocytes also did not alter tumor expansion in the peritoneal cavity (Fig. 3g).

Altogether, we conclude that the lack of mature adipocytes in the omentum or mesentery does not grossly impact the types of immune cells that accumulate at these sites. It is also unlikely, based on the minimal impact of broad lymphocyte and NK cell depletion on tumor expansion in the peritoneal cavity, that the omentum’s primary role in governing OC tumor seeding and growth in the peritoneum is linked to the modulation of activities of lymphocytes or NK cells.

### Metabolic syndrome does not exacerbate OC growth in the peritoneum

While overall tumor burden, organ distribution, and spatial localization were equivalent in PARC and DARL mice whose metabolic parameters were normalized with fat pad transplants, systemic metabolic dysregulation or metabolic syndrome may alter omental tumor seeding patterns. To evaluate this possibility, Adiponectin^Cre+^ DTA^fl/+^ lipoatrophic mice with metabolic syndrome (Extended Data Fig. 2) not corrected by fat transplant were injected i.p. with ID8p53^−/−^Brca2^−/−^GFP Luc tumor cells. Tumor growth was monitored and compared with that of female littermate controls lacking Cre expression, which consequently exhibited normal fat distribution. Despite elevated blood triglyceride and metabolic perturbations (Extended Data Fig. 2), Adiponectin^Cre+^ DTA^fl/+^ lipoatrophic mice had a similar overall tumor burden compared to control littermates (Fig. 4a). Omentectomy reduced the pace of tumor growth in lipoatrophic mice and control mice, even in the context of metabolic syndrome (Fig. 4b). The omentum still comprised 50-60% of the overall tumor burden in control and lipoatrophic mice (Fig. 4c). Approximately 80% of the omental fat region consisted of tumor nodules in both lipoatrophic and control omenta. In comparison, tumors occupied less than 10% of the corresponding adjacent mesothelial sheets. Similarly, about 30% of the mesenteric fat region was occupied by tumors in both groups, with less than 5% of the adjacent mesothelial sheet occupied (Fig. 4d).

**Figure 4.**
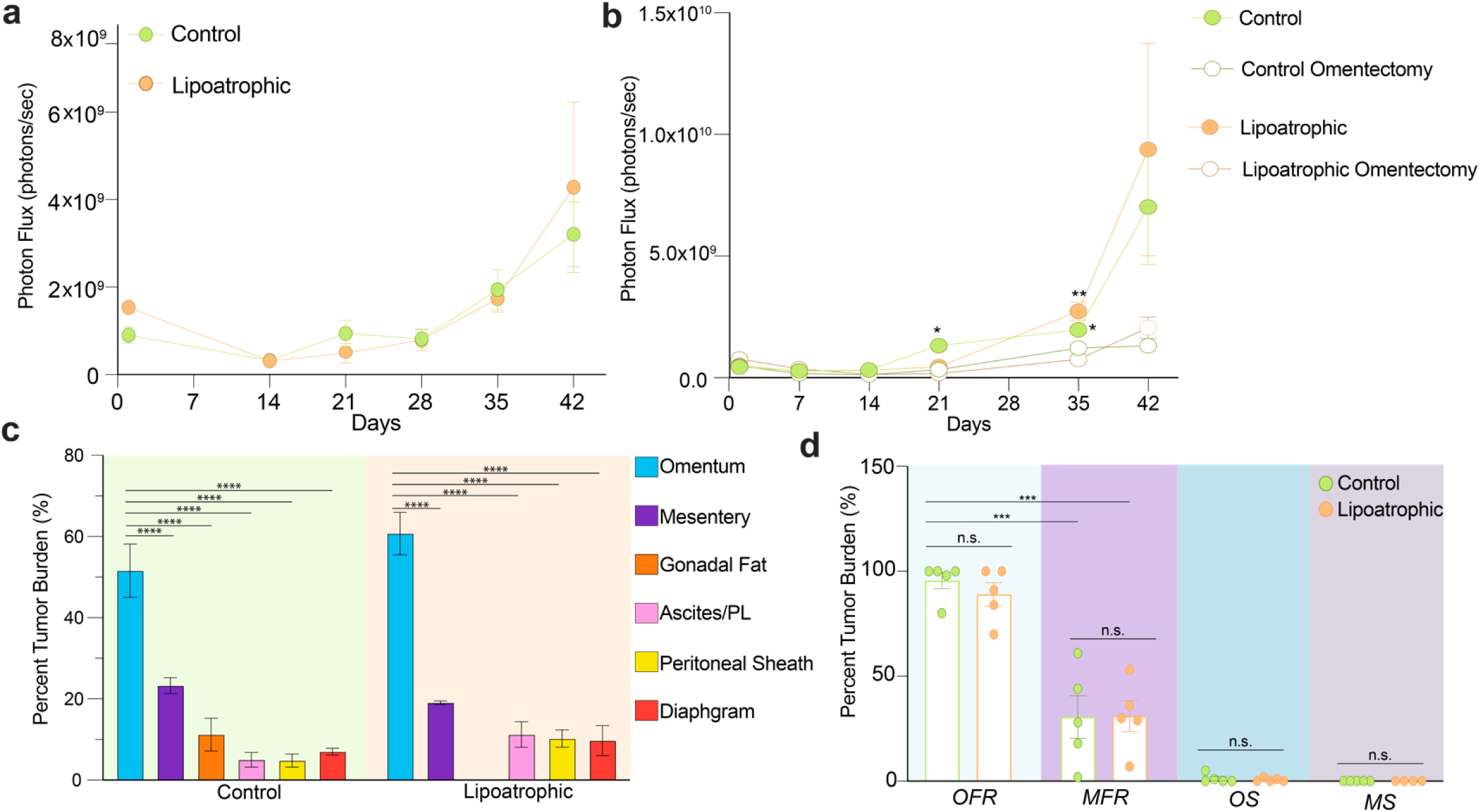
Metabolic syndrome and lipoatrophy do not mediate OC omental tropism in the peritoneal cavity. **a**. Quantification of in vivo bioluminescence images of control (*n*=4) and lipoatrophic (*n*=4) mice post i.p. injection of 5×10^6^ ID8p53^−/−^Brca2^−/−^ GFP Luc tumor cells. A two-way ANOVA (n.s. for genotype) and post-hoc comparisons using Tukey’s HSD were conducted to compare the means of the groups over time, n.s at any time point. **b**. In vivo bioluminescence of control and lipoatrophic mice that received either a sham surgery (control, *n*=10; lipoatrophic, *n*=7) or omentectomy (control, *n*=13; lipoatrophic, *n*=7) followed by an i.p. injection of 5×10^6^ ID8p53^−/−^Brca2^−/−^ GFP Luc tumor cells six weeks later. Two-way ANOVA and post-hoc comparisons using Tukey’s HSD were conducted to compare the means of the groups over time, n.s at any time point between control and lipoatrophic mice and n.s. at any time point between control omentectomized and lipoatrophic omentectomized mice. Omentectomized control mice compared to littermates *P<0.05 at days 21 and 35. Omentectomized lipoatrophic mice compared to littermates **P<0.05 at day 35. c. Distribution of tumor burden across peritoneal organs based on quantifying ex vivo bioluminescence images of pinned organs in control (*n*=4) and lipoatrophic (*n*=4) mice six weeks after i.p. injection of 5×10^6^ ID8p53^−/−^Brca2^−/−^ GFP Luc tumor cells. Two-tailed Student t-test, ***P<0.001, ****P<0.001 **d**. Percent of the omentum fat region (OFR), mesentery fat region (MFR), omentum sheet (OS), and mesenteric sheet (MS) occupied by tumor in control (*n*=4) and lipoatrophy (*n*=4) mice six weeks post 5×10^6^ ID8p53^−/−^Brca2^−/−^ GFP Luc tumor injection using measurements from FIJI Image J particle analysis. Two-Tailed Student’s t-test between control and lipoatrophic OFR, OS, MFR, MS, n.s. Two-Tailed Student’s t-test between lipoatrophic or control OFR and MFR, ***P<0.001.

We conclude that hypertriglyceridemia and aberrant glucose tolerance, which characterize lipoatrophic mice and elements of clinical obesity, do not alter the peritoneal adhesion or expansion of OC tumors, nor do these metabolic parameters affect the critical role of the omentum in driving such growth.

### PARC and DARL mice have similar cell composition and high FABP4 endothelial expression in the omentum compared to the mesentery

We ruled out a major anticipated role for mature adipocytes or systemic metabolic changes such as hypertriglyceridemia or glucose intolerance in driving OC expansion in the peritoneal cavity. At the same time, we found the omentum was highly linked to tumor growth in all settings. Thus, we set out to identify features of the omentum that might explain its key role and reconcile other known findings with the present results. We conducted comprehensive gene expression profiling to compare the cell populations and gene expression profiles of the omentum and mesentery in lipoatrophic mice with those of their control littermates. The sheet and fat regions of the omentum and mesentery from 8 PARC and DARL mice were disaggregated to generate cell suspensions for single-cell RNA sequencing. CD45^+^ and CD45^-^ cells were sorted separately, combined, and submitted for single-cell RNA sequencing to allow for unbiased assessment of any population, immune or stromal, that might differentially shift between the PARC and DARL mice peritoneal organs (Extended Data Fig. 4b). The resulting dataset included 69,807 cells classified in 18 Louvain clusters, of which 25,617 were *Cd45^-^*and 44,190 were *Cd45*^+^. Hashtags or antibody-derived tags (ADTs) were used in a process termed ‘Cell Hashing’ to label eight biological replicates for each tissue independently^58^. This hashtag ADT labeling was effective, as association with 1 of each of the eight hashtags collectively accounted for 62.2% of the data when the standard hashtag demultiplexing analysis was performed (see Methods). The difference between the two most abundant ADT hashtags in each cell had a median difference of 630 cells, enabling the accurate assignment of one ADT hashtag per cell (Extended Data Fig. 6a) and the identification of biological replicates across each tissue. The remaining data were identified as multiplets (20.7%) or were negative for any confident hashtag signal (17.1%; Extended Data Fig. 5b, c). Reasonable biological variance was observed across replicates for tissues and cell types (Extended Data Fig. 6d-f).

Data from the omenta (sheet and adipose region) and mesentery (sheet and adipose region) of PARC and DARL mice were clustered based on differentially expressed genes in each cluster using canonical markers for specific populations (Fig. 5a). Overall, there were few differences in the clustering of cells between PARC and DARL mice (Fig. 5b). Since mature adipocytes were anticipated to be scarce in the dataset due to their failure to pellet during centrifugation or to be tagged in the 10X sequencing pipeline, the lack of mature adipocytes in even the PARC samples was expected. Furthermore, the proportions of different *Cd45^+^*cell types and identifiable stromal cell clusters across the PARC and DARL mice were comparable (Fig. 5c). Because not all *Cd45*^-^ stromal cell types could be categorized using canonical markers, Capybara, a computational single-cell classifier, was used to assign the stromal cells’ identities relative to selected published references^59^. In brief, we used the computational tool Capybara to compare our data with biologically similar scRNA-seq datasets for which cell type classification has already been performed (see Methods). Capybara provides utility in cell type classification for datasets with no identical predecessors in the published literature, such as this one. Relative similarities and approximations can be measured across cell types present in each reference and query dataset. Across multiple reference comparisons^43,45,60^, there were no meaningful proportional shifts in any identified stromal cell population between the PARC and DARL tissues (Fig. 5d), including pre-adipocyte and adipocyte progenitor populations, that may stimulate matrix remodeling to support OC metastasis^61^. We anticipated that pre-adipocyte populations would be amplified in DARL mice, given the absence of mature adipocytes that express adiponectin; however, the pre-adipocyte populations were similar between the cohorts. Differential gene expression, based on fold changes greater than 5-fold, revealed that Retnla and Cfd (complement factor D) were enriched in PARC over DARL cells, serving as a strong positive control and validation for our model, considering that these genes are well-established adipokines^62,63^ (Extended Data Fig. 6g, Table 1).

**Figure 5.**
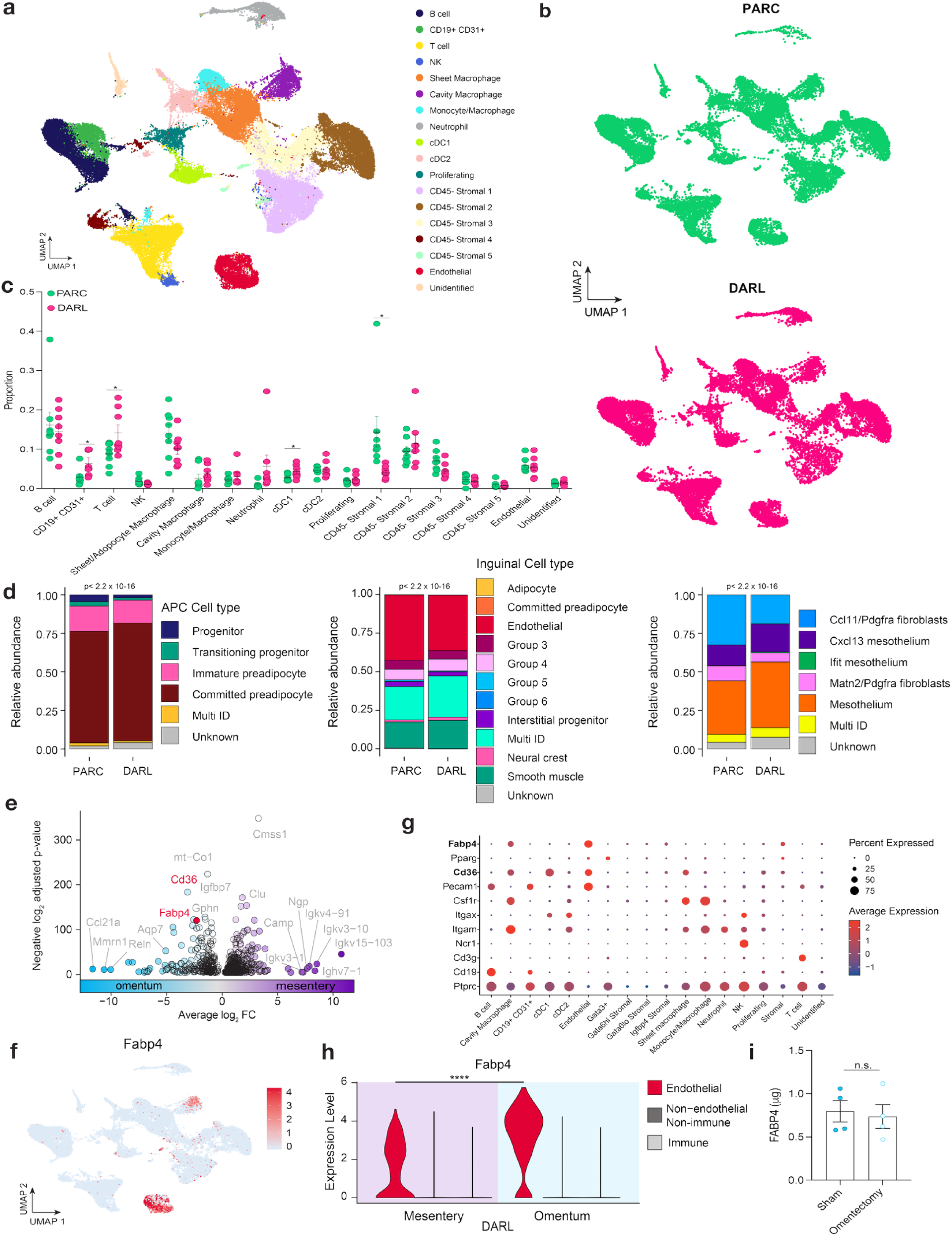
PARC and DARL mice share transcriptional profiles with enriched FABP4 in omental endothelial cells. **a**. UMAP visualization of all cells collected from the omentum and mesentery sheet and fat regions in PARC (*n*=8) and DARL (*n*=8) mice. **b**. UMAP visualization of all PARC (top) and DARL (bottom) cells separately. **c**. Comparison of the number of cells in each identified cluster between PARC and DARL mice. Two-Tailed Student’s t-Test, *P<0.05. **d**. Cabybara classification of Cd45^-^ stromal cells using three different classification schemes based on existing cell identity assignments. Left, middle, and right plots were generated from Capybara comparisons to mouse skin Lin-/Cd29+/Cd34+ APC cells isolated from SVF of naive mice ^43^, Mouse inguinal CD45^-^ APC cells isolated from SVF of naive mice (age P12)^60^, and mouse omental CD45^-^/CD41^-^/Ter119^-^/CD31/PDPN^+/−^ stromal cells from naive mice (age 8-12 weeks)^45^, respectively. Chi-Square Test was performed on the capybara distribution analyses for each reference to determine if the cell type distribution varied between genotypes (α=0.05). **e**. Differential gene expression in the omentum (blue) and mesentery (purple) identifies mRNAs, including *Fabp4* mRNA, that were more highly expressed in the omentum than in the mesentery. **f**. UMAP visualization of FABP4 expression across all cells from the omentum and mesentery sheet and fat regions in PARC (*n*=8) and DARL (*n*=8) mice. **g**. Dot plot of specific genes expressed in clustered cell types. *Fabp4* is highly expressed in endothelial cells and to a lesser extent in macrophages. **h**. Violin plot illustrating FABP4 expression is highest in the endothelial cells compared to other immune and non-immune subtypes. Unpaired t-test with Welch’s correction, two-tailed, ****P<0.0001. **i.** Quantification of circulating FABP4 in plasma collected from sham or omentectomized mice (*n*=4). Two-Tailed Student’s t-Test, n.s.

Once we established that the immune and stromal composition of PARC and DARL mice were similar, we compared differentially expressed genes in the omentum and mesentery that might distinguish the tumor kinetics and proliferation in these organs (Fig. 5e). *Fabp4* expression was significantly more dominant in the omentum than in the mesentery (Fig. 5e). FABP4 was demonstrated to be pivotal in OC tumor expansion in the peritoneal activity. This observation fueled the concept that adipocyte-derived lipids were critical participants in tumor growth^9,64,65^. Yet, our data indicated that mature adipocytes are not needed to drive tumor growth. To reconcile these data, we wondered which cell types in the omentum other than mature adipocytes might be significant sources of FABP4. Two clusters in our dataset, cavity macrophages and endothelial cells, showed high expression of *Fabp4* mRNA (Fig. 5a, f, g, h). However, *Fabp4* mRNA expression was far more pronounced in the endothelium than in macrophages (Fig. 5g). The endothelial cluster was further differentiated and classified to identify arterial, venous, capillary, and lymphatic endothelial cells using known markers and the Capybara algorithm to match a known endothelial reference^66^ (Extended Data Fig. 6 i,j). Most of the endothelial cells in the dataset were blood vascular rather than lymphatic endothelium (Extended Data Fig. 6 i,j). *Fabp4* was broadly expressed in the endothelium but was exceptionally high in endothelial cells with lower levels of *Cxcl12* (Extended Data Fig. 6j). Notably, distinguishing the omentum from the mesentery, including in DARL mice that preserved tumor growth in the absence of mature adipocytes, *Fabp4* expression in omental endothelium was markedly elevated over that in the endothelium of the mesentery or immune cells of either tissue (Fig. 6h). Thus, the expression of *Fabp4* as a known pivotal player in OC growth in the peritoneum that was previously attributed to adipocytes is robustly and differentially elevated in endothelial cells of the omentum, the critical site in the peritoneal cavity that supports the most robust tumor expansion regardless of its affiliation with mature adipocytes, lymphocytes, or NK cells. These findings help reconcile previous evidence that omental FABP4 contributes to ovarian cancer spread by identifying endothelial cells as a key source, independent of adipocytes. The evidence that circulating FABP4 arises from endothelial cells^33^, combined with the high expression of Fabp4 in omental endothelial cells in our single-cell sequencing data, we wondered if the omentum may support a significant proportion of circulating FABP4. However, when we assessed this possibility by ELISA, there was no difference in plasma FABP4 levels between mice that received sham surgery and those that received omentectomy (Figure 5i). Thus, while omentectomized mice have decreased tumor burden (Figure 2j), it is unlikely that changes in circulating FABP4 can explain this difference. However, endothelial FABP4 may have a local effect on tumor burden.

**Figure 6.**
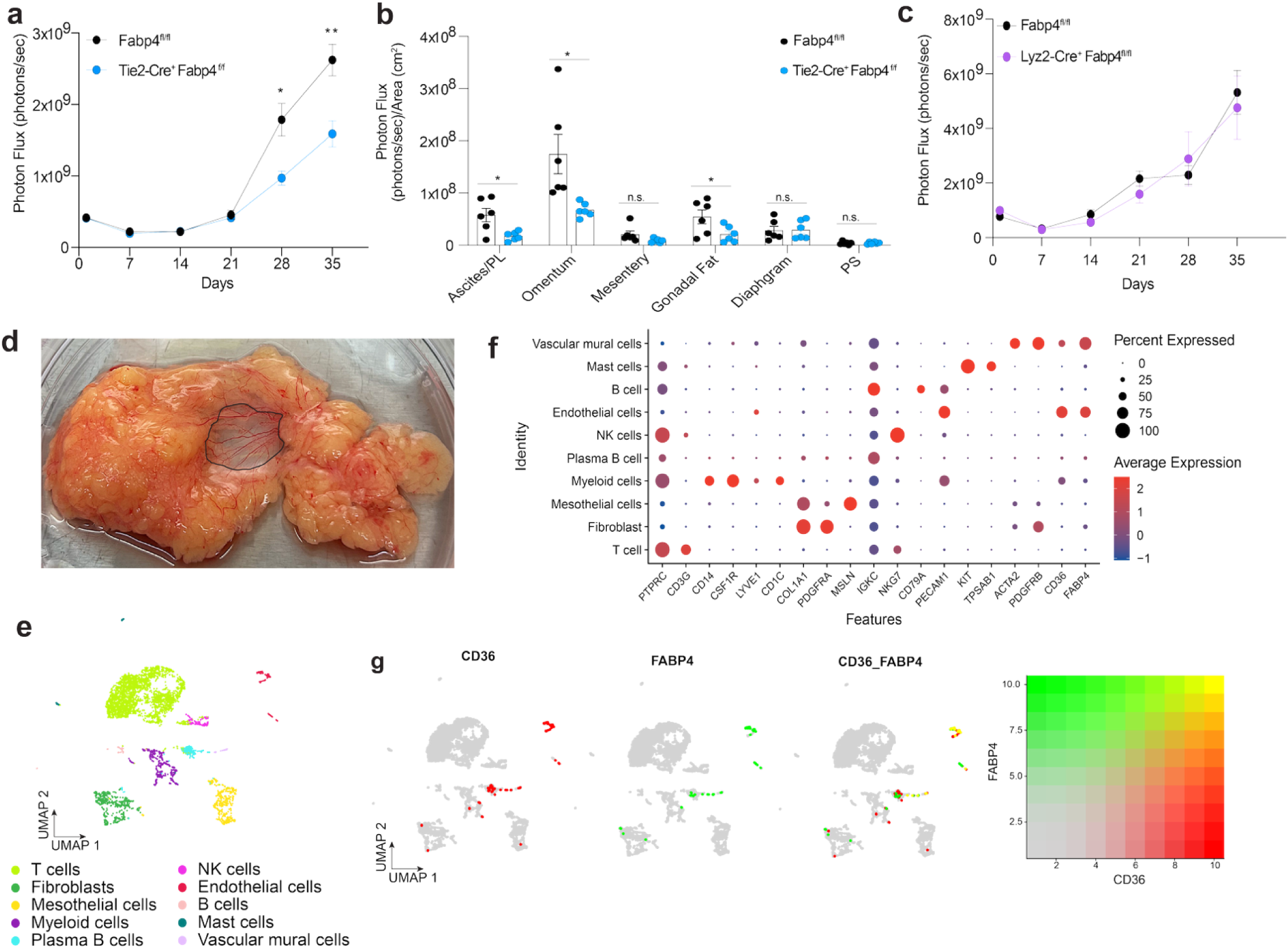
Endothelial FABP4 promotes omental tumor burden and defines a conserved FABP4⁺CD36⁺ vascular population. **a**. Quantification of in vivo bioluminescence images of Tie2-Cre⁺ FABP4^fl/fl^ mice and littermate controls **b**. Distribution of tumor burden across peritoneal organs based on quantifying ex vivo bioluminescence images of pinned organs in Tie2-Cre⁺ FABP4^fl/fl^ mice and littermate controls (n=6/group) at day 35. PS = peritoneal sheath**. c.** Quantification of in vivo bioluminescence images of Lyz2-Cre⁺ FABP4^fl/fl^ mice and littermate controls (n=6/group) **d**. Benign human omentum tissue obtained for single-cell RNA sequencing. The translucent sheet (outlined in black) and adipocyte-rich regions were dissected and sequenced separately for spatial specificity. **e**. UMAP projection of all cells from human omental single-cell RNA-seq, showing 11 transcriptionally defined clusters. **f**. Dot plot of specific genes expressed in clustered cell types. *FABP4 is* highly expressed in endothelial cells and vascular mural cells. **g**. Feature plots and co-expression analysis reveal that FABP4 and CD36 are co-expressed in a subset of human omental endothelial cells, suggesting a conserved lipid-handling vascular phenotype. Data represent mean ± SEM. *p < 0.05, **p < 0.01 by two-way ANOVA or unpaired t-test, as appropriate.

### Endothelial FABP4 promotes omental tumor burden and defines a conserved FABP4⁺CD36⁺ vascular population

To test the role of FABP4 expressed in endothelium in impacting the growth of OC in the peritoneal cavity, we bred endothelial-specific FABP4 knockout mice (Tie2-Cre⁺ Fabp4^fl/fl^) along with littermate controls^33^. Following i.p. injection of ID8p53^−/−^Brca2^−/−^ GFP Luc tumor cells, tumor burden and expansion lagged in mice lacking endothelial FABP4 (Figure 6a). Ex vivo BLI further confirmed a marked reduction in tumor signal localized to the omentum (Figure 6b), suggesting that FABP4-expressing endothelial cells play a role in supporting omental tumor establishment or growth.

Because FABP4 was also expressed in macrophages in the murine omentum (Figure 5g) and Tie2 can be induced in some hematopoietic cells^67^, we tested whether myeloid-lineage FABP4 played a comparable role. Lyz2-Cre⁺ Fabp4^fl/fl^ mice were injected i.p. with ID8p53⁻/⁻ Brca2⁻/⁻ GFP-Luc tumor cells. In contrast to the endothelial cell deletion model, no significant difference in tumor burden was observed (Figure 6c), indicating that the tumor-promoting function of FABP4 within the endothelium is not shared by immune cell populations such as macrophages.

To assess the relevance of FABP4-expressing omental endothelial cells in humans, we obtained benign human omental tissue from the Washington University OB/GYN tissue bank. The translucent sheet region, outlined in black, and the adjacent adipocyte-rich region were carefully microdissected, enzymatically digested, and processed as separate samples for single-cell RNA sequencing. Unsupervised clustering of the adipocyte region identified 11 distinct cell populations (Figure 6e). FABP4 expression was enriched in endothelial cells, which also expressed PECAM1, and in vascular mural cells that expressed contractile or smooth muscle cell-associated genes, such as ACTA2 and 2RB (Figure 6f). Notably, CD36, a lipid transporter, was co-expressed in the same cluster. Blended feature plots confirmed co-expression of FABP4 and CD36 (Figure 6g), underscoring the presence of a FABP4⁺CD36⁺ endothelial cell population in the human omentum that may have a prominent lipid-transporting role. Thus, endothelial cells of the omentum in both mice and humans express elevated mRNA encoding FABP4 and CD36. FABP4 within these cells promotes tumor growth in the omentum. When these data are combined with the evidence that we provided here that omental adipocytes do not affect the pace of OC growth in the omentum or peritoneal cavity, it seems reasonable to conclude that earlier implications that FABP4 promoted ovarian tumor growth in omentum may relate to its key role in the vasculature, rather than the adipocyte.

## Discussion

OC is known for organ tropism, particularly to the omentum and other peritoneal surfaces, as per Paget’s original “seed and soil” hypothesis^68^. However, more nuanced seeding patterns exist within organs such as the omentum. The canonical omentum is the adipose-rich region embedded in well-vascularized connective tissue covered by mesothelial lining continuously resting on the basement membrane, excluding where the milky spots form. In addition to the adipose region, the omentum includes a thin, net-like, fenestrated, translucent sheet composed of mesothelial cell layers that are poorly vascularized and enclose diffuse collagen fibers^3^. Some studies suggest that mesothelial cells support tumor growth^69^. Other studies suggest that neutrophil extracellular traps^70^, omental macrophages^71,72^, blood vessels^73^, fibroblasts^74^, or lymphoid aggregates called milky spots^75–79^, consisting of immune cells such as B cells and macrophages, support tumor growth. Furthermore, adipocytes, the cells that define the omentum as a fat pad, have also been argued to drive OC metastasis to the omentum due to their ability to support tumor fatty acid metabolism^80,81^. The roles of these different cell types or tissue components are not mutually exclusive.

Our data show that tumors not only gravitate to peritoneal organs of adiposity, such as the omentum and mesentery, but also selectively engraft in these adipocyte-rich regions rather than in the tissue sheet area of tissue lacking adipose tissue. Nonetheless, unexpectedly, the mature adipocytes that demarcate these spaces were not responsible for tumor engraftment. Although adipocytes were absent in critical fat pads such as the omentum in lipoatrophic mice, cancer still preferentially grew where the adipocytes would have been localized. These findings suggest that a critical niche exists where adipocytes are typically found that is conducive to cancer implantation and growth. However, this niche can support cancer progression even in the absence of mature adipocytes, indicating that they are dispensable. Furthermore, the omentectomized lipoatrophic mice have decreased tumor burden, indicating that functionally, the omentum still maintains a dominant role in governing tumor burden in the absence of adipocytes. The lack of adipocytes also did not alter the response to carboplatin therapy.

Because the omentum contains unique lymphoid aggregates called milky spots, the immune cells could contribute to tumor localization in the omentum. Our flow cytometry analysis revealed changes in the immune populations in the omentum and mesentery at steady state. Nonetheless, neither antibody depletion nor genetic knockout models, which led to the loss of T cells, B cells, and NK cells, resulted in increased tumor burden in the omentum, running counter to other tumor models where the role of immune surveillance was first recognized through greater tumor growth in lymphocyte-deficient mouse strains^82^. This result is also in accordance with findings in multiple fallopian tube-derived HGSOC cell lines, such as the KPCA and BPPNM lines used in some experiments herein, in which monotherapy with anti–PD–1 antibodies had minimal impact on tumor progression^39^. Together, these results suggest that while T cells may be present in the OC milieu, they may not be sufficiently primed, recruited, or functionally capable of mediating anti-tumor activity. This interpretation aligns with clinical trial data showing limited survival benefit from immune checkpoint inhibitors, either as monotherapy or in combination, in HGSOC^83^. While adipocytes are often proposed to exert immunosuppressive effects that may contribute to this resistance^21,84^, our data argue against this being a dominant mechanism, as even in mice lacking mature adipocytes, adaptive immune cell depletion did not significantly impact tumor progression.

Based on the equivalent tumor phenotype between control and lipoatrophic mice, it is unsurprising that single-cell RNA sequencing did not reveal major differences in immune or stromal populations between the omentum and mesentery, regardless of whether mature adipocytes were present. While FABP4-expressing mature adipocytes have been proposed as critical mediators of OC progression^9^, those findings relied on whole-body knockout models, complicating interpretation due to FABP4’s expression in multiple non-adipocyte compartments, including endothelium and macrophages^85,86^. Our murine single-cell RNA-seq analysis revealed that, at the tissue level, FABP4 expression was higher in the omentum than in the mesentery. Within the omentum, endothelial cells were the primary source of FABP4 expression, both in the presence and absence of mature adipocytes. This finding indicates that elevated endothelial FABP4 is an intrinsic feature of the omental microenvironment, rather than a consequence of local adiposity. The identification of a conserved FABP4^+^CD36^+^ endothelial population in human omentum further supports the clinical relevance of this vascular specialization in establishing a tumor-permissive niche. These data suggest that endothelial, rather than adipocyte-derived, FABP4 may be a key determinant of the omentum’s unique tumor-supportive niche. Circulating FABP4 has also emerged as a key regulator of metabolism^33,87,88^. However, omentectomy, which effectively decreases the overall tumor burden, did not result in changes in FABP4 levels in the plasma. This result suggests that the role of FABP4 expressed by the endothelium functions in a local, rather than systemic context, when it comes to impacting OC expansion in the omentum.

Endothelial FABP4 may instead play a role in omental tumorigenesis by regulating angiogenesis. Given that adipose-rich regions of the omentum and mesentery are highly vascularized, it is plausible that tumors preferentially localize to these sites due to their vascular density. Notably, FABP4 is regulated downstream of VEGFA through DLL4-NOTCH1 signaling, a pathway well known for regulating endothelial tip-stalk dynamics^86^. Supporting this, prior studies using chitosan-based nanoparticles to deliver siRNA specifically to tumor vasculature demonstrated that silencing endothelial FABP4 reduced angiogenesis and peritoneal tumor spread in OC models^89,90^. Extending these findings, we show that genetic deletion of FABP4 in endothelial cells (Tie2-Cre⁺ Fabp4^fl/fl^)^33^ significantly reduces omental tumor burden. In contrast, deletion of FABP4 in myeloid cells had no measurable impact on tumor progression, demonstrating the cell-type specificity of FABP4’s function in the tumor microenvironment. As an alternative to considering a role for endothelial FABP4 in angiogenesis, it may function to facilitate fatty acid shuttling to tumor cells through its role as a fatty acid transporter. While adipocytes have long been implicated in providing fatty acids to fuel tumor cells, our data suggest that vascular cells themselves may serve as central metabolic contributors, particularly through lipid-handling programs such as FABP4.

The identification of FABP4⁺CD36⁺ endothelial cells in both murine and human omentum supports a shift in the understanding of peritoneal metastasis from an adipocyte-mediated to an endothelium-mediated metabolic mode. This finding aligns with the growing recognition that endothelial cells play active, cell-specific roles in lipid handling, which influence the tissue and tumor microenvironments. For example, CD36, a fatty acid transporter traditionally associated with adipocyte-induced expression in OC cells^91^, is also expressed in endothelial cells, where it facilitates the uptake of long-chain fatty acids and contributes to vascular lipid metabolism^92–94^. CD36 enables fatty acid uptake, which is crucial for metastasis-initiating cells, and CD36 inhibition blocks metastasis without affecting primary tumor growth^95^. Similarly, the lipoprotein receptor SR-BI plays an endothelial-specific role in LDL transcytosis and atherosclerosis, independent of its hepatic function^96^, suggesting that endothelial lipid transporters can have distinct, tissue-specific functions relevant beyond metabolic disease. Caveolin-1 (CAV1), the core structural protein of caveolae—lipid-rich membrane invaginations involved in lipid transport and signaling—demonstrates cell-type–specific functions as well. Its endothelial deletion disrupts lipid uptake^97^, impairs angiogenesis^98^, and differs functionally from adipocyte CAV1 in metabolic regulation^99^. These cases underscore the importance of contextualizing lipid-handling programs by cell type, supporting our conclusion that endothelial FABP4 is a conserved and functionally specific contributor to the omental metastatic niche.

While the notion that adipocytes drive OC growth has long dominated the field, our study reframes this paradigm by functionally uncoupling the role of mature adipocytes from peritoneal tumor progression. By preserving systemic metabolic homeostasis while eliminating mature adipocytes, we demonstrate that adipocytes are dispensable for omental tumor localization, expansion, and chemotherapy response. Instead, our findings position the endothelium, specifically a conserved FABP4⁺CD36⁺ population in the omentum, as a metabolically active and tumor-supportive compartment. This vascular specialization challenges the adipocyte-centric model and underscores the importance of considering stromal and metabolic features of the omental microenvironment in shaping peritoneal metastasis.

## Data Availability

Murine single-cell sequencing data generated in this study have been deposited in the Gene Expression Omnibus (GEO) (RRID:SCR_005012) under accession code GSE281706. Human single-cell sequencing data generated in this study have been deposited in the GEO (RRID:SCR_005012) under accession code GSE300038. Publicly available single-cell RNA sequencing datasets can be found in references^42–44^ and under GEO accession numbers GSE140348, GSE128889, GSE 241627. All other data supporting the findings of this study are available from the corresponding author upon request.

## Acknowledgements.

Major support for this work included NIH grant R37AI049653 to GJR, NIH F30CA251124 to RLM, NIH grant K00CA264434 to JH, and NIH grant R01 CA258325 to WZ. SAM is now affiliated with the Division of Gastroenterology, Hepatology and Endoscopy, Division of Genetics, Brigham and Women’s Hospital; Department of Systems Biology, Harvard Medical School, Boston, Massachusetts. We thank the Genome Technology Access Center at the McDonnell Genome Institute at Washington University School of Medicine for help with genomic analysis. The Center is partially supported by NCI Cancer Center Support Grant #P30 CA91842 to the Siteman Cancer Center from the National Center for Research Resources (NCRR), a component of the National Institutes of Health (NIH), and NIH Roadmap for Medical Research. We thank Julie Prior from the Molecular Imaging Center at Washington University School of Medicine for allowing us to use the IVIS50 for bioluminescence imaging studies. The Center is supported by NCI P30 CA091842 (Siteman Cancer Center Small Animal Cancer Imaging shared resource), NIH-S10OD027042, and NNIH S10D025264. This publication is solely the authors’ responsibility and does not necessarily represent the official view of NCRR or NIH. We thank the Washington University Gynecology Oncology Tissue Bank for providing human omentum samples. This work was done in partial fulfillment of the PhD studies requirements for RLM. Many thanks to the members of her thesis committee—Dineo Khabele, Katherine Weilbaecher, Patricia Ribeiro Pereira, and Quing Zhu—for their invaluable project advice and support. We thank Beth Helmink for their surgical training in mice and overall advice. We are grateful to Robert Schreiber and members of his laboratory, including Hussein Saltan, for providing advice and antibodies for lymphocyte depletion. We thank Karen Inouye and Gökhan S. Hotamişligil (Harvard) for providing us with mice to use and for Karen’s feedback on the manuscript draft. Lastly, we thank Emma Erlich and Raphael Czepielewski for consistent feedback, technical advice, and experimental guidance.

## Author contributions

RLM, GJR conceptualized and planned the study; EGB, JH, AG, SN, AK, conducted experiments and analyzed data; MW and NKHY established the mouse model in the lab; NZ, KEI, and GSH generated and provided key tools; WZ provided surgical assistance with experiments; SAM, BHZ, GJR provided supervision and obtained regulatory compliance and funding; RLM and GJR drafted the manuscript with editing input from all authors.

