## Supplemental figures for "Adipocytes are dispensable in shaping the ovarian cancer tumor microenvironment in the omentum"

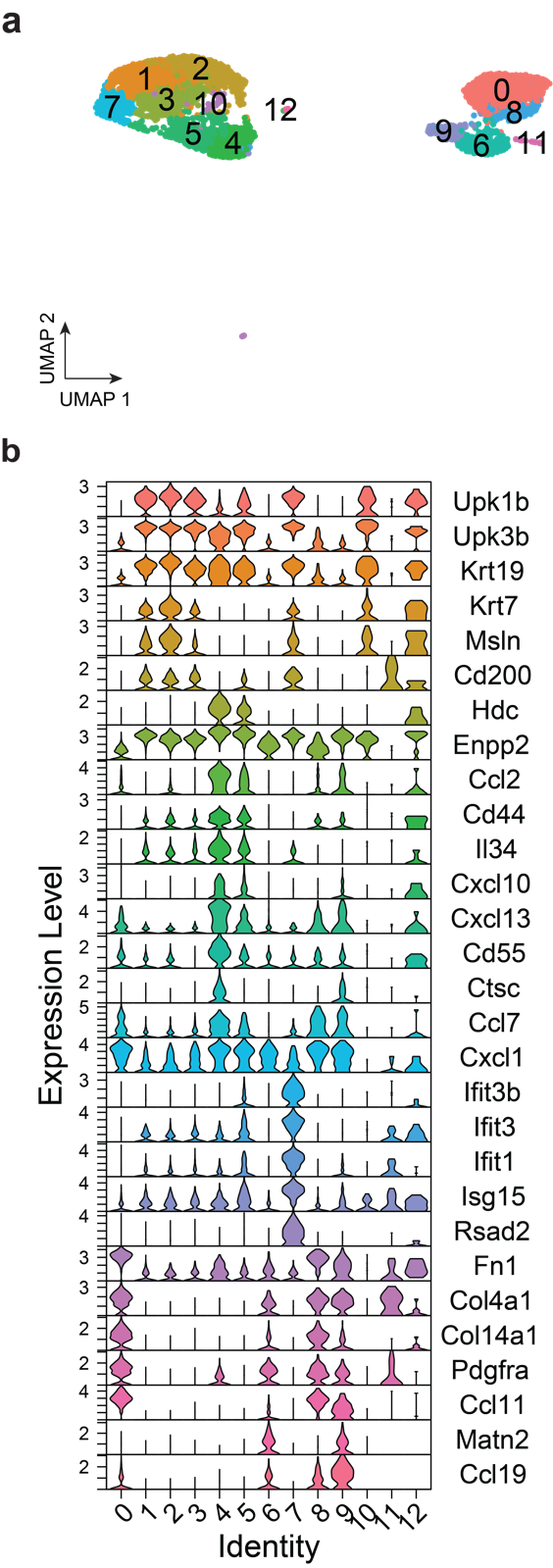

Supplemental Methods 1. Cell type classification of Jackson-Jones *et al*. **a**. UMAP embedding of non-endothelial stromal cells from the omentum, colored by Louvain clusters. For data published by Jackson-Jones *et al*., raw data were obtained from GEO and re-processed (GSM4053741;^1^). 3,817 were used to construct the reference, following the removal of cells with fewer than 1,000 genes per cell (*nFeature_RNA*). Louvain clustering was performed using the default parameters of the Seurat function *FindClusters()* (Seurat package version 5.1.0). **b**. Violin plots of expression of marker genes used to identify cell types as described by the authors. Louvain clusters 1, 2, 3, 4, 5, 7, 10, and 12 were identified as mesothelial by positive expression of epithelial and mesothelial markers *Upk1b*, *Upk3b*, *Krt19*, *Krt7*, *Msln*, and *Cd200*. A subset of these, clusters 4 and 5, were identified as Cxcl13+ mesothelium by positive expression of *Hdc*, *Enpp2*, *Ccl2*, *Cd44*, *Il34*, *Cxcl10*, *Cxcl13*, *Cd55*, *Ctsc*, *Ccl7*, and *Cxcl1*. Another subset of these, cluster 7, was identified as Ifit+ mesothelium by positive expression of *Ifit3b*, *Ifit3*, *Ifit1*, *Isg15*, and *Rsad2*. Louvain clusters 0, 8, and 9 were identified as Ccl11+/Pdgfra+ fibroblasts by positive expression of *Fn1*, *Col4a1*, *Col14a1*, and *Pdgfra*. Louvain cluster 6 was identified as Matn2+/Pdgfra+ fibroblasts by positive expression of *Matn2* and *Ccl19*. Finally, Louvain clusters 11 and 12 were not assigned any cluster described by the authors.

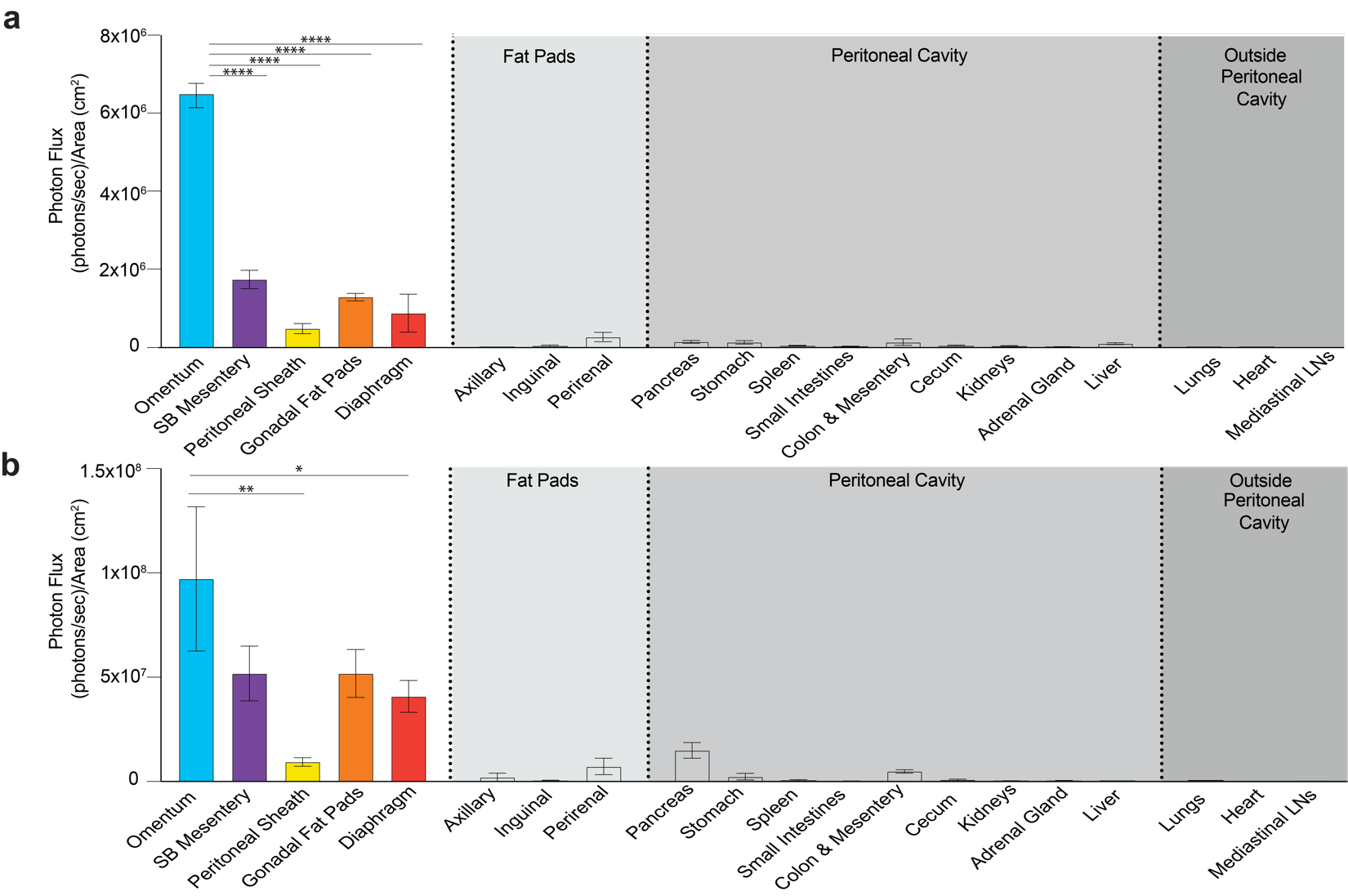

Extended Data Figure 1. Biodistribution of tumor burden in C57BL/6J WT mice (*n*=5) at **a**. one-week (One-way ANOVA****P<0.0001) and **b**. six weeks (One-way ANOVA *P<0.05) post i.p. injection of 5x10^6^ ID8p53^–/–^Brca2^–/–^ GFP Luc tumor cells based on quantifying the photon flux in ex vivo bioluminescence images. Tukey’s HSD Test, ****P<0.0001, **P<0.01, *P<0.05. SB= small bowel; LN = lymph node
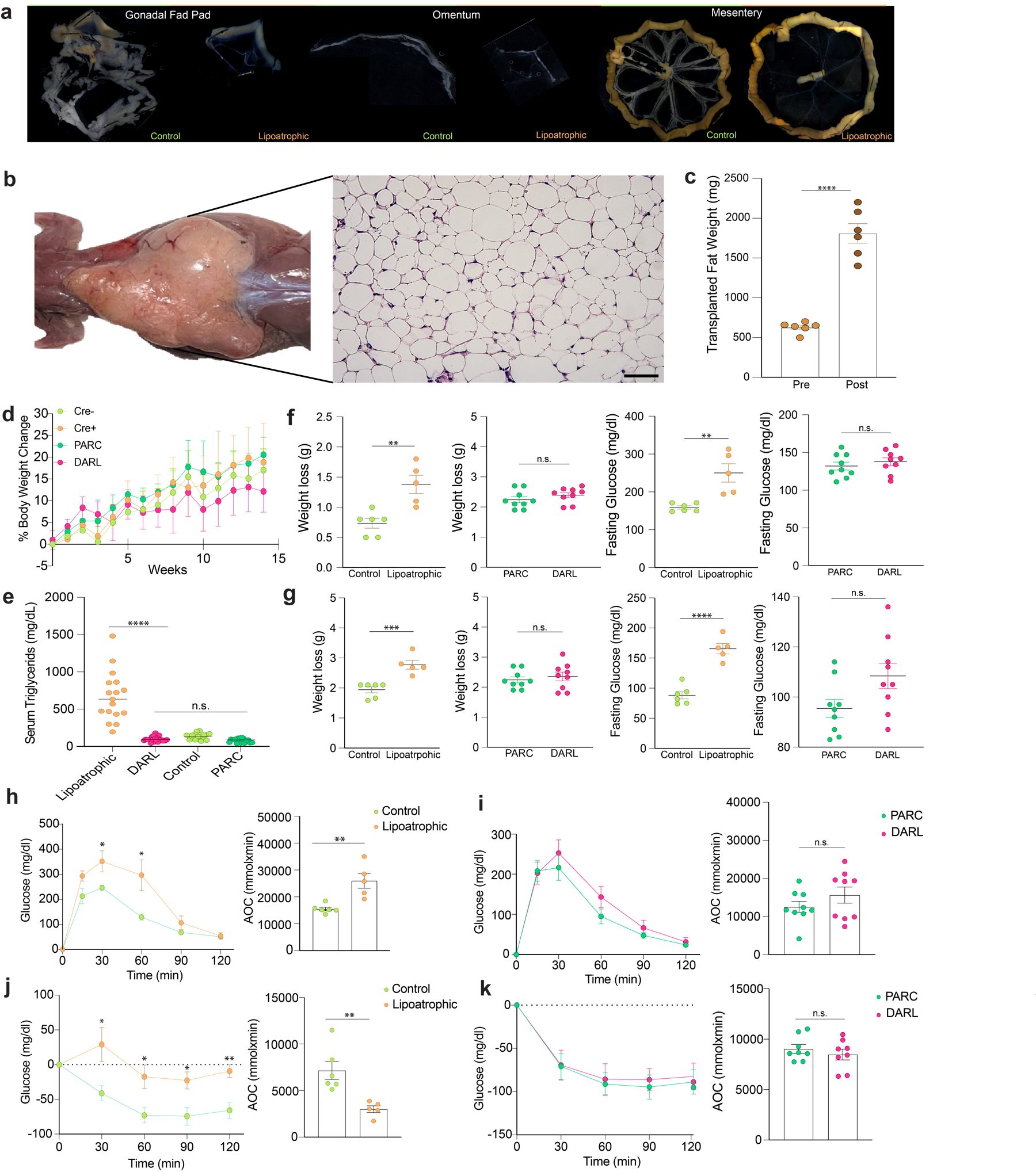

Extended Data Figure 2. Fat pad transplant surgeries normalize metabolic parameters in lipoatrophic mice. **a**. Representative stereoscope brightfield images of peritoneal fat regions in control and lipoatrophic mice, including the gonadal fat pads, omentum, and mesentery. **b**. Brightfield stereoscope and corresponding H&E image of a fat transplant in a lipoatrophic mouse 12 weeks post fat transplant surgery (Scale bar, 50µm) **c**. Weight of transplanted fat pre and 12 weeks post-transplant in a lipoatrophic mouse. Two-Tailed Student’s T-test, ****P<0.0001. **d**. Weight of control (*n*=15) and lipoatrophic (*n*=6) mice without surgery and corresponding PARC (*n*=9) and DARL (*n*=8) littermates that received fat transplant surgery normalized to the weight at the time of surgery. Two-way ANOVA and post-hoc comparisons using Tukey’s HSD were conducted to compare the means of the four groups over time, n.s. at any time point. **e**. Serum triglyceride measurements in control (*n*=18) and lipoatrophic (*n*=17) mice pre-surgy and PARC (*n*=17) and DARL (*n*=20) mice post fat transplant surgery. One-way ANOVA and post-hoc comparisons using Tukey’s HSD, ****P<0.0001. Weight loss and fasting glucose levels in control (*n*=5) and lipoatrophic mice *(n*=5) and corresponding PARC (*n*=9) and DARL (*n*=9) mice after fasting for **f**. 6 or **g**. 12 h. Two-tailed Student’s t-test **P<0.01, ***P<0.001, ****P<0.0001. Glucose tolerance test in **h**. Control (*n*=8) and lipoatrophic (*n*=8) mice before surgery and i. in PARC (*n*=8) and DARL (*n*=8) mice post fat transplant surgery. AOC= area of the curve. Two-Tailed Student’s t-test at each time point, *P<0.05, **P<0.01. Insulin tolerance test in **j**. Control (*n*=8) and lipoatrophic (*n*=8) mice before surgery, and **k**. PARC (*n*=8) and DARL (*n*=8) mice post fat transplant surgery. AOC= area of the curve. Two-Tailed Student’s t-test at each time point, *P<0.05, *P<0.01

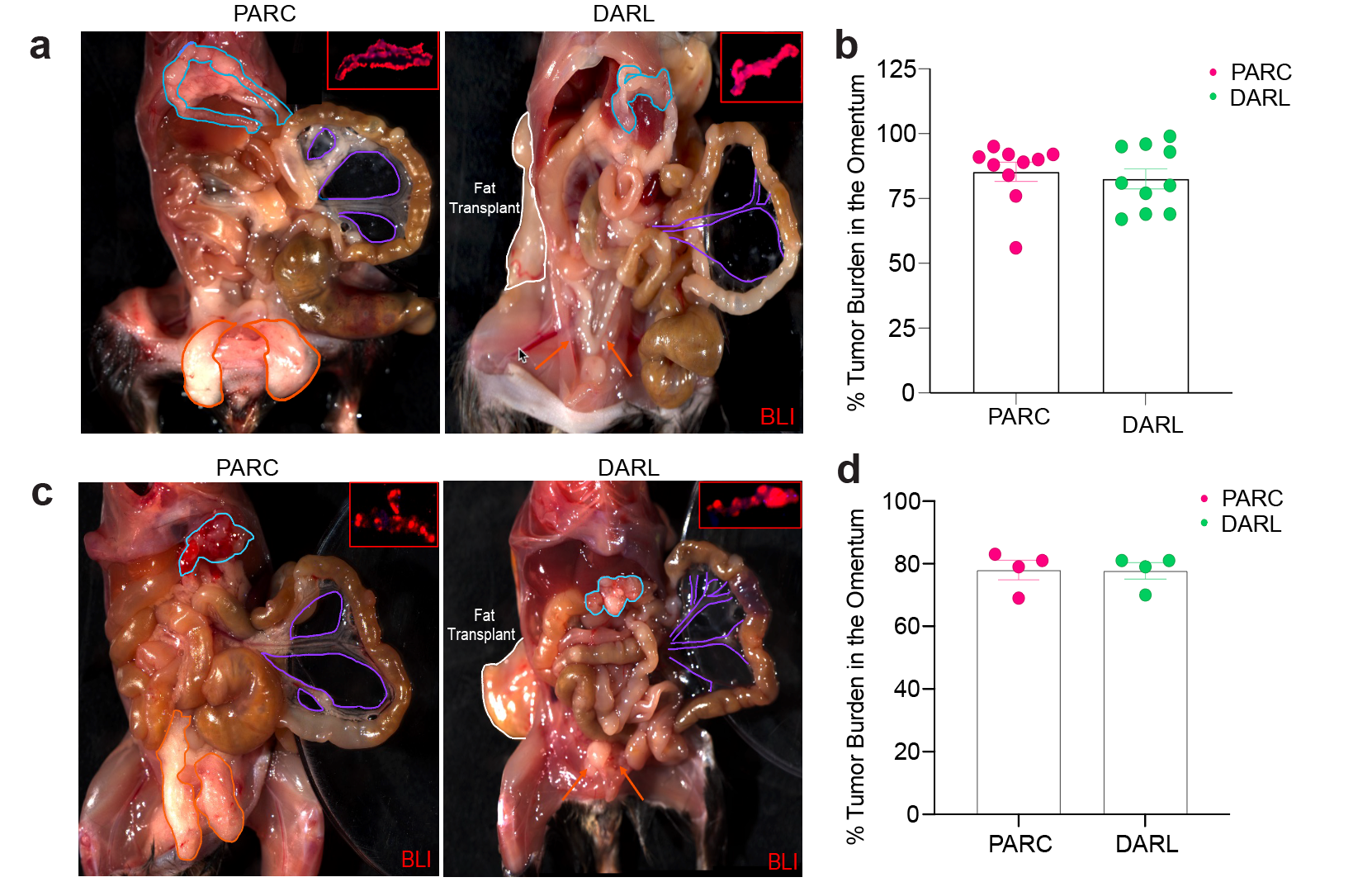

Extended Data Figure 3. Omentum tumor niche is maintained even in the absence of mature adipocytes. Representative images and ex vivo tumor quantification in the omentum of tumor-injected PARC and DARL mice at endpoint following i.p. injection of 1x10^6^ KPCA-luc tumor cells (**a.**, **b.**) or 3x10^6^ BPPNM-luc tumor cells **(c.**, **d.**) are depicted. Harvest of KPCA-injected mice occurred at day 7; BPPNM-injected mice were harvested at day 21. Panels a and c show tumor encapsulation of the omentum in both PARC and DARL groups. Blue outlines the omentum. Purple outlines the mesenteric fat region. Orange outlines the gonadal fat, or the orange arrows point to the gonadal fat region in DARL mice. The white outlines of fat transplants in DARL mice. The inlet in the top right is the ex vivo bioluminescence photo of the omentum, illustrating bioluminescent tumor nodules in red. Quantification of the percent omental tumor burden (b, d) demonstrates comparable localization to the omentum regardless of adipocyte presence.

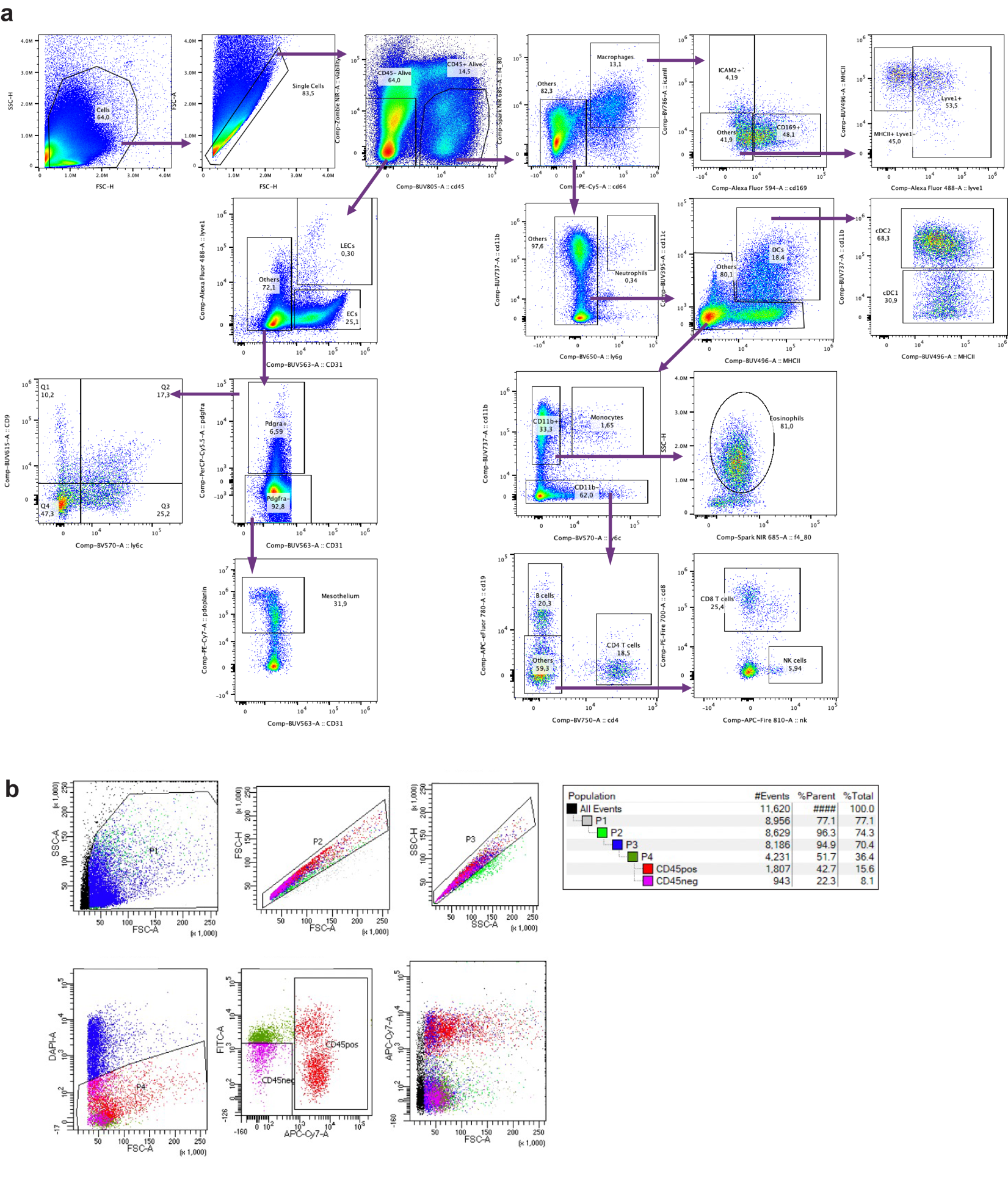

Extended Data Figure 4. Flow cytometry and cell sorting strategies. **a**. Flow cytometry gating to classify CD45+ immune cell subsets in PARC (*n*=4) and DARL (*n*=4) mice at steady state. **b**. Sorting scheme to collect PARC and DARL CD45+ and CD45- populations from omentum and mesentery samples (sheet region or fat region) for single-cell RNA sequencing.

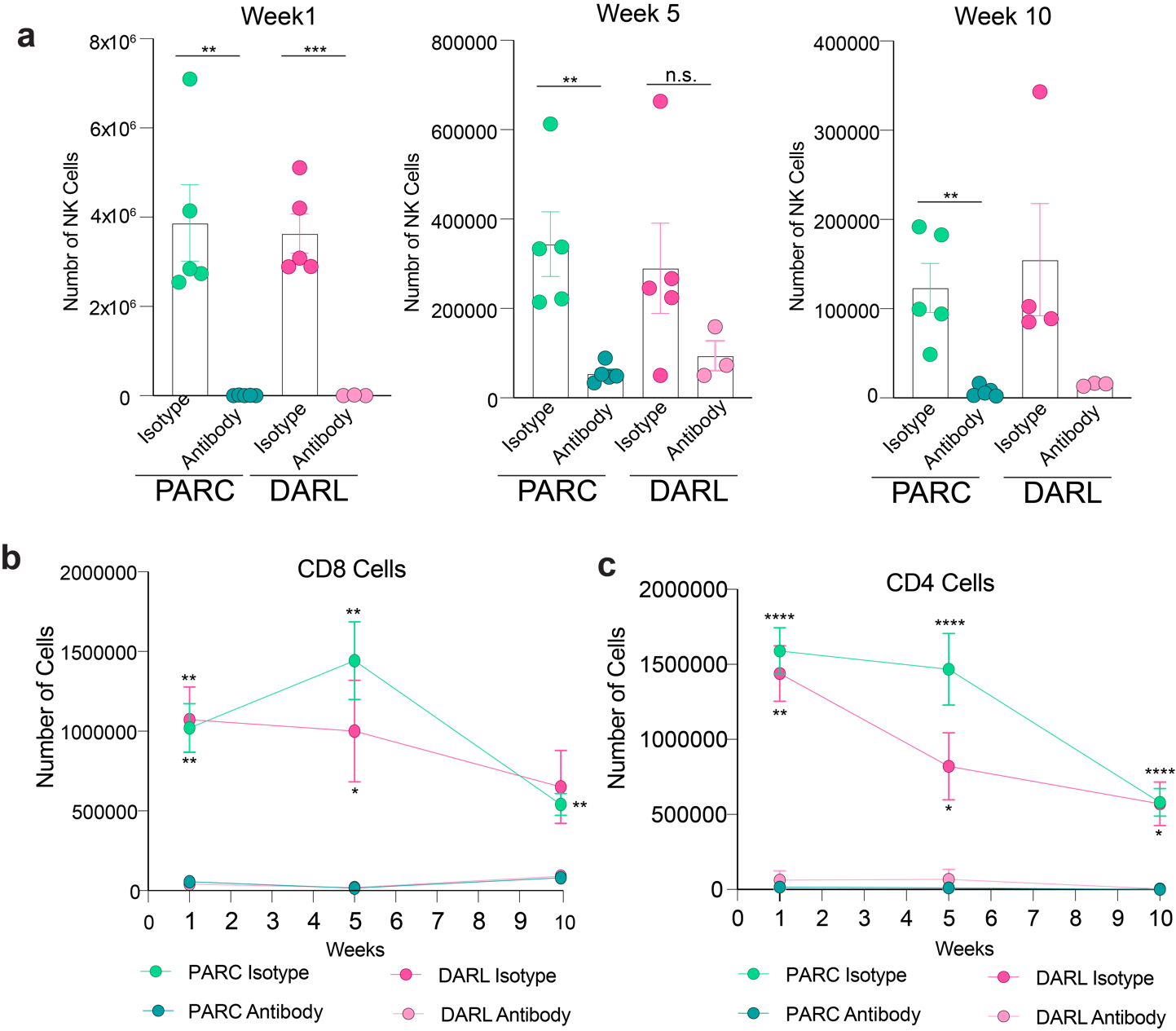

Extended Data Figure 5. Validation of in vivo immune cell depletion. **a**. NK (Natural Killer) **b**. CD8^+^ and **c**. CD4^+^ cell numbers over time in PARC (*n*=5) and DARL mice (*n*=5) given N1.1, CD4, and CD8 depleting antibodies (*n*=3) or matched isotype control antibodies (*n*=5). Two-Tailed Student t-test comparing DARC isotype and DARC antibody treated mice or PARC isotype and PARC antibody treated mice, *P<0.05, **P<0.01, ***P<0.001, ****P<0.0001.
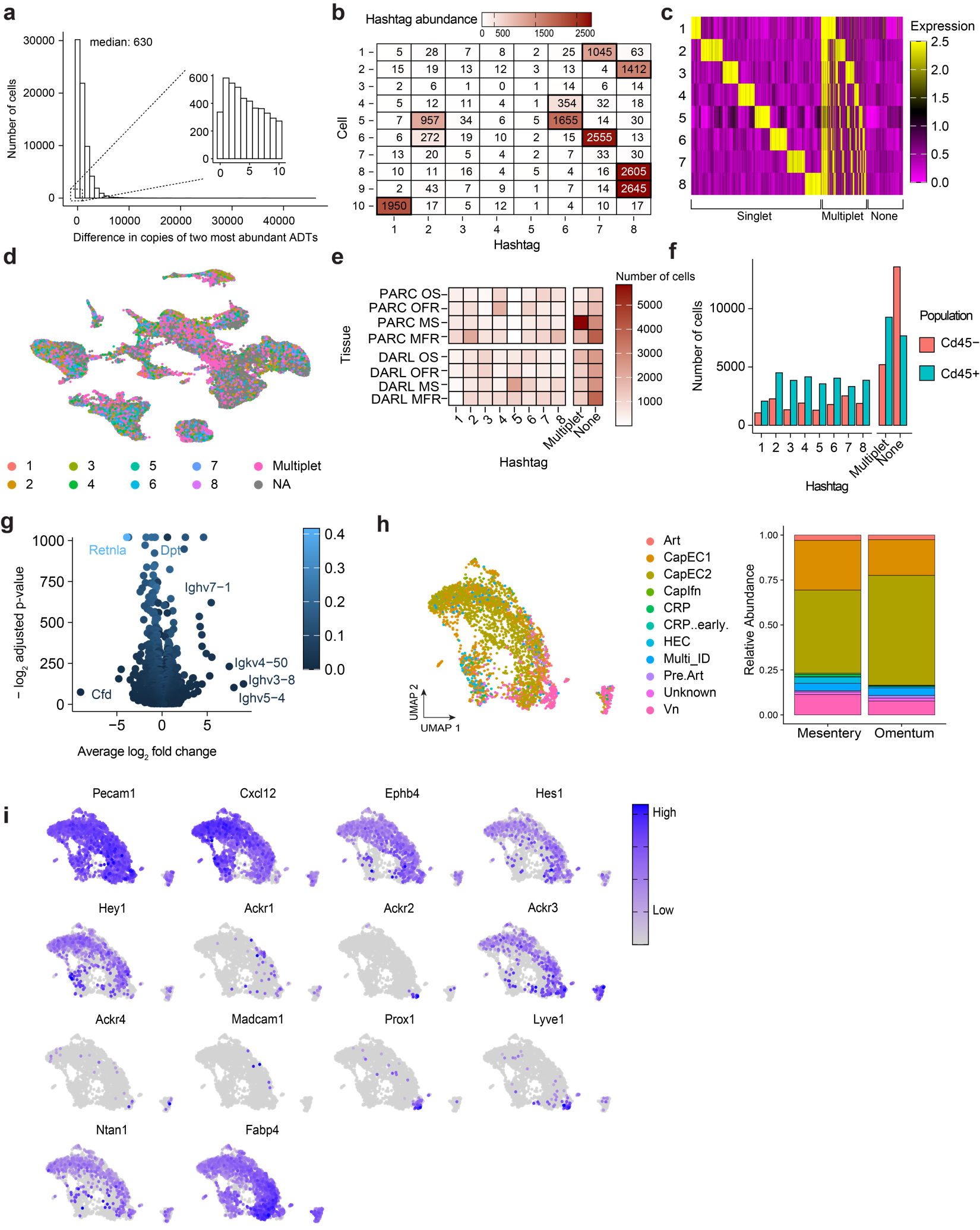

Extended Data Figure 6. Antibody derived tags or hashtag analysis and endothelial characterization of PARC and DARL single-cell RNA sequencing data. **a**. Histogram showing the difference in copies between the two most abundant antibody-derived tags (ADTs) in each cell across all PARC (*n*=8) and DARL (*n*=8) tissues, including the omentum fat region (OFR), mesentery fat region (MFR), omentum sheet (OS), and mesenteric sheet (MS). **b**. Representative table of the hashtag assignment for 10 randomized cells based on the number of copies of the oligo found in each cell. Bolded tiles denote hashtag assignment for each cell. Cells 3 and 7 were not assigned to any hashtag; cells 5 and 6 were assigned as multiplets of hashtags 2/6 and 2/7, respectively. **c**. Heat map of scaled (z-scores) normalized antibody counts sorted by classification for a random sample of 5000 cells. **d**. UMAP visualization of the hashtag distribution across all cells. **e**. Quantification of the cell numbers across the eight hashtag assignments in all tissues. **f**. Quantification of *Cd45+* and *Cd45-* cells across all PARC and DARL tissues (OFR, OS, MFR, MS). **g**. Dot plot of differentially expressed markers between *Cd45*-/DARL and *Cd45*-/PARC cells after Bonferroni correction of p-values. Features with positive fold change are upregulated among DARL cells; features with negative fold change are upregulated among PARC cells. Features are colored by the absolute difference in the fraction of cells expressing each feature across the two groups. **h**. UMAP visualization of the Capybara comparison to cell identities classified in a murine lymph node blood vascular endothelium dataset.^2^ (left). Comparison of the endothelial cell types identified by Capbybara between the omentum and mesentery endothelium (right). **i**. Co-localization of *Fabp4* expression with endothelial cluster sub-classifications based on the expression of canonical genes.

Extended Data References

1	Jackson-Jones, L. H. *et al.* Stromal Cells Covering Omental Fat-Associated Lymphoid Clusters Trigger Formation of Neutrophil Aggregates to Capture Peritoneal Contaminants. *Immunity* **52**, 700-715.e706 (2020). <https://doi.org/10.1016/j.immuni.2020.03.011>

2	Brulois, K. *et al.* A molecular map of murine lymph node blood vascular endothelium at single cell resolution. *Nature Communications* **11**, 3798 (2020). <https://doi.org/10.1038/s41467-020-17291-5>
